# Clustered PHD domains in KMT2/MLL proteins are attracted by H3K4me3 and H3 acetylation-rich active promoters and enhancers

**DOI:** 10.1101/2021.10.15.462366

**Authors:** Anna Maria Stroynowska-Czerwinska, Magdalena Klimczak, Michal Pastor, Asgar Abbas Kazrani, Katarzyna Misztal, Matthias Bochtler

## Abstract

Histone lysine-specfic methyltransferase 2 (KMT2A-D) proteins, alternatively called mixed lineage leukaemia (MLL1-4) proteins, mediate positive transcriptional memory. As the catalytic subunits of human COMPASS-like complexes, they methylate H3K4 at promoters and enhancers. KMT2A-D contain understudied highly conserved triplets and a quartet of plant homeodomains (PHDs). Here, we show that all clustered PHDs localise to the well-defined loci of H3K4me3 and H3 acetylation-rich active promoters and enhancers. Surprisingly, we observe little difference in binding pattern between PHDs from promoter-specific KMT2A-B and enhancer-specific KMT2C-D. Fusion of the KMT2A CXXC domain to the PHDs drastically enhances their preference for promoters over enhancers. Hence, the presence of CXXC domains in KMT2A-B, but not KMT2C-D, may explain the promoter/enhancer preferences of the full-length proteins. Importantly, targets of PHDs overlap with KMT2A targets and are enriched in genes involved in the cancer pathways. We also observe that PHDs of KMT2A-D are mutated in cancer, especially within conserved folding motifs (Cys4HisCys2Cys/His), which cause a domain loss-of-function. Taken together, our data suggests that PHDs of KMT2A-D guide the full-length proteins to active promoters and enhancers, and thus play a role in positive transcriptional memory.

**GRAPHICAL ABSTRACT:** 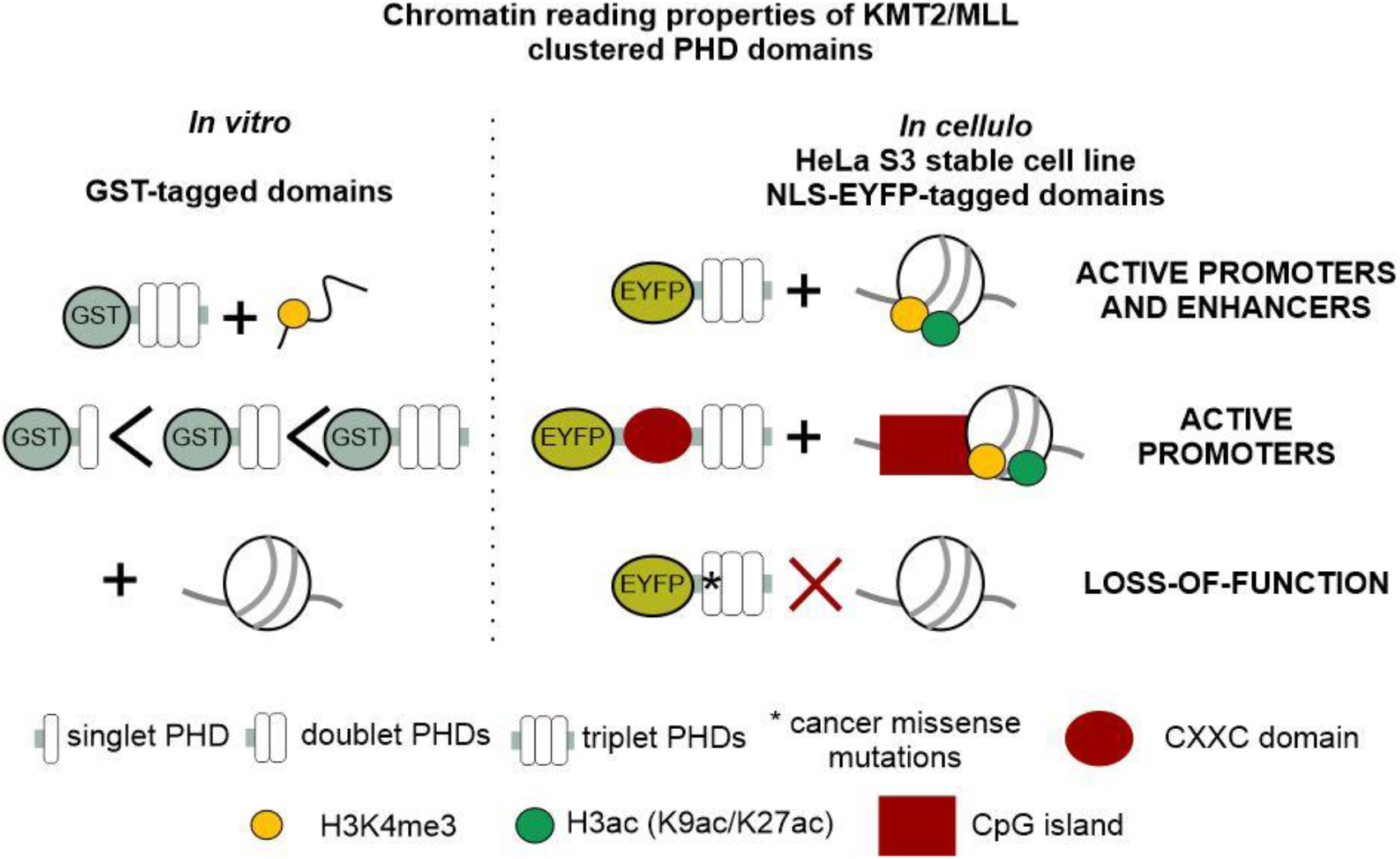

## INTRODUCTION

Histone lysine-specific methyltransferases KMT2A-D are catalytic engines of human COMPASS-like (hCOMPASS-like) complexes that mediate positive transcriptional memory, by depositing the activating mark H3K4me1-3 at regulatory elements (promoters and enhancers) of already active genes [1–4]. hCOMPASS-like complexes share many subunits, including WD repeat-containing protein 5 (WDR5), Retinoblastoma-binding protein 5 (RbBP5), Absent small homeotic-2 like (ASH2L), and Dumpy-30 (DPY-30) that together form the shared WRAD sub-complex [5]. KMT2A-D are alternatively termed as mixed lineage leukaemia proteins (MLL1-4), as a result of KMT2A/MLL1 originally being identified as a frequent breakpoint in mixed lineage leukaemia (MLL) [6]. KMT2C/MLL3 and KMT2D/MLL4 mutations are also found in other malignancies [7,8] and frequent enough that KMT2C/MLL3 features prominently in lists of pan-cancer drivers [9–11].

hCOMPASS-like complexes containing KMT2A-B are recruited to active promoters, where they deposit the active promoter modification H3K4me3 [4,12–14]. In contrast, the hCOMPASS-like complexes with KMT2C or KMT2D preferentially bind to enhancers, where they deposit the enhancer mark H3K4me1 [15–17]. The complex chromatin targeting is influenced by both non-catalytic subunits [14,18,19] and intrinsic preferences of the KMT2A-D proteins [20]. However, the mechanistic causes for the preference for promoters or enhancers is not well understood.

The domain organization of the KMT2A-D proteins provide some clues about their binding preferences (Fig. 1a). Only promoter-specific KMT2A and KMT2B possess CXXC domains which bind non-modified CpGs [21–23], the hallmark of active promoters [24]. Furthermore, all KMT2A-D proteins contain several chromatin readers, the PHD domains (PHDs). Stand-alone PHDs are widely present in transcription regulating proteins from plants to animals, and even in yeast [25]. These compact domains consist of around 65 amino acids and contain a Cys4HisCys2Cys/His (C4HC2C/H) folding motif, which chelates two structural Zn^2+^ ions [25]. PHD domains are known primarily as readers of H3K4me3 [26,27], which they accommodate in a deep, hydrophobic pocket that is present in many PHD domains [28]. In addition, rare double PHD domains in MOZ and DPF bind acetylated lysine (H3K14ac) in a deep, hydrophobic pocket, which is distinct from the methyl binding pocket [29,30].

**Figure 1.**
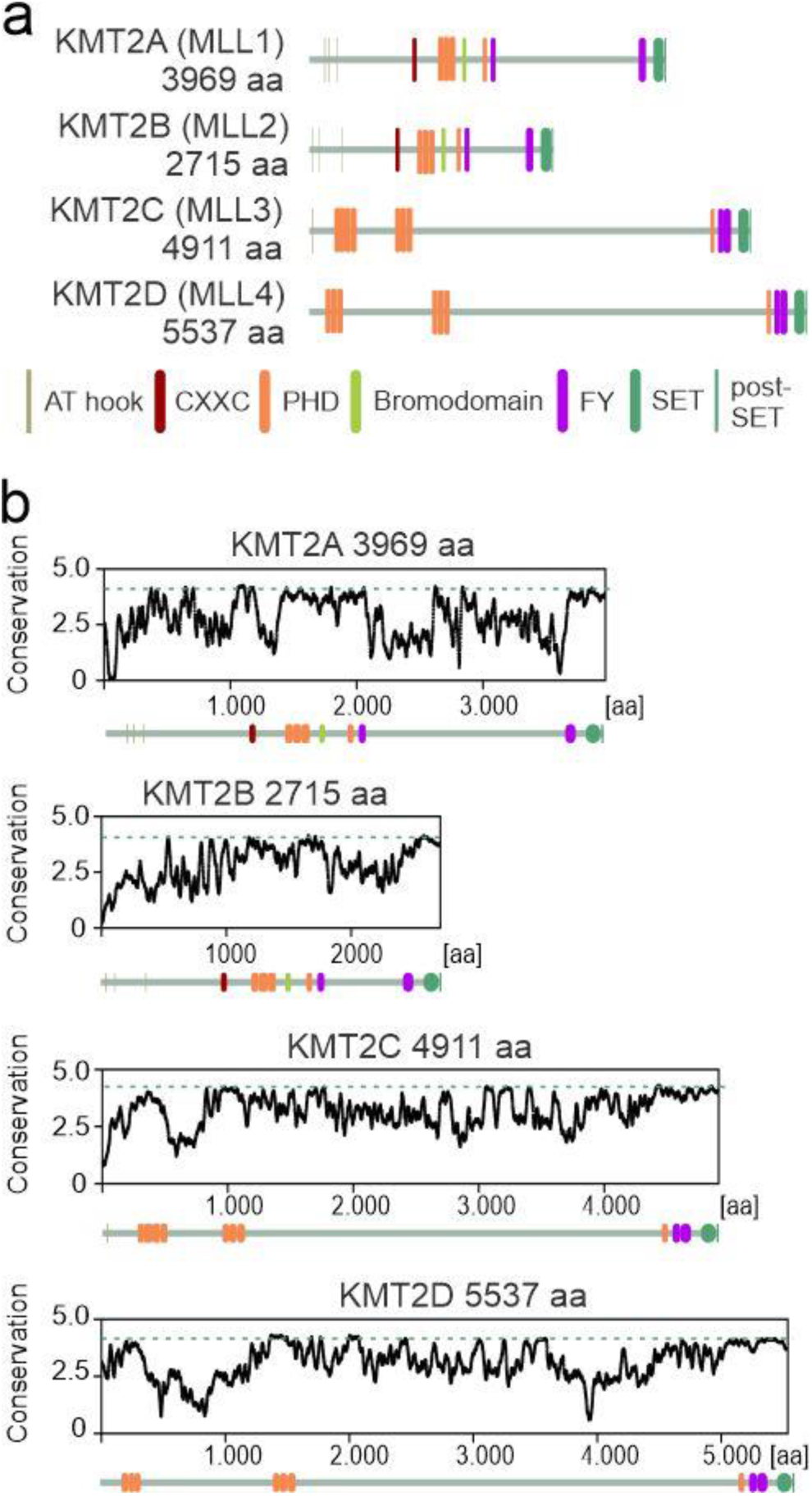
Clustered PHD domains (PHDs) are highly conserved in KMT2A-D proteins. **a** KMT2A-D proteins architecture. **b** Conservation of KMT2A-D proteins across species, measured in bits of information as in sequence logos. Domain architecture is indicated below the graphs. All diagrams are on the same scale. The dashed line shows the extent of conservation of the SET domain.

In KMT2A-D, most PHD domains are arranged in triplets or even a quartet. This arrangement is unique in the human proteome but has received only little attention. Prior studies of PHD domains in KMT2A-D have typically focused on single PHD domains and a set of candidate chromatin marks as potential binding partners (H3K4me3, non-modified H4, H4R3me2, H4K16ac, H4K20ac and H4K20me1/3) [28,31–33]. Surprisingly, H4K16ac have been found to be bound by PHD6 of KMT2D outside the deep acetylation pockets, in a fairly shallow region of the protein surface [31].

In this research, we aimed to understand the promoter and enhancer specificity of KMT2A-D proteins by characterizing PHD domains with respect to their chromatin binding properties, specifically in the context of cellular chromatin environment. We present a first unbiased genome-wide characterization of the cellular binding sites of clustered PHDs in KMT2A, KMT2C and KMT2D proteins. We show that all the clustered PHDs bind preferentially to well-defined loci in acetylation-rich active promoters and enhancers (rich in H3K9ac, H3K27ac but also H3K4me3) of cancer-related genes. The loci bound by PHDs overlap partially but significantly with targets of full-length hCOMPASS-like subunits (KMT2A, WDR5 and RbBP5). Fusion of the CXXC domain to the clustered PHDs of KMT2A increases its affinity to promoters and improves recapitulation of binding properties of the full-length protein. Our experimental data and the frequent loss-of-function cancer mutations in KMT2A-D PHD domains point to the important role of PHD domains in targeting hCOMPASS-like complexes to active promoters and enhancers.

## RESULTS

### PHDs are highly conserved in KMT2A-D proteins

Amino acid conservation between PHDs of some mammals is surprisingly high (~99% identity) and prompted us to analyse conservation of orthologues from a broader range of species (Suppl. Fig. 1a). This comparison indicated that PHDs are among the most conserved regions within KMT2A-D proteins (Fig. 1b), in some cases (i.e., PHD4-6 of KMT2D) rivalling conservation of the catalytic SET domains. Differences between paralogous clustered PHDs in human KMT2A-D proteins are considerable, with typical values for amino acid identity and similarity below 40% and 50%, respectively (Suppl. Fig. 1b). The high conservation between species and variability among different modules suggests that PHDs play an important role in KMT2A-D biology.

### Interaction between PHDs and histones strengthens with an increasing number of PHDs

For *in vitro* biochemical characterisation, we cloned and purified from *E. coli* bacterial lysates all clustered PHDs of KMT2A-D proteins as well as some singlets or doublets as fusion proteins with GST-tag (Fig. 2a-c). We excluded PHDs from KMT2B from further studies, as they were poorly expressed and had low solubility. Hereafter, we designate individual PHDs by their paralogue identifier (2A-D) and by their position, i.e., the third PHD in KMT2A is abbreviated as 2A3. Similarly, clusters of PHDs are designated by the first and last domains, i.e., 2A13 (comprising PHD1, PHD2 and PHD3 of KMT2A). PHDs containing an extension with the C4 zinc finger are termed “extended” PHDs [32] and are abbreviated with an extra “e”: 2Ae3, 2Be3, 2Ce4, 2Ce7, and 2De6.

**Figure 2.**
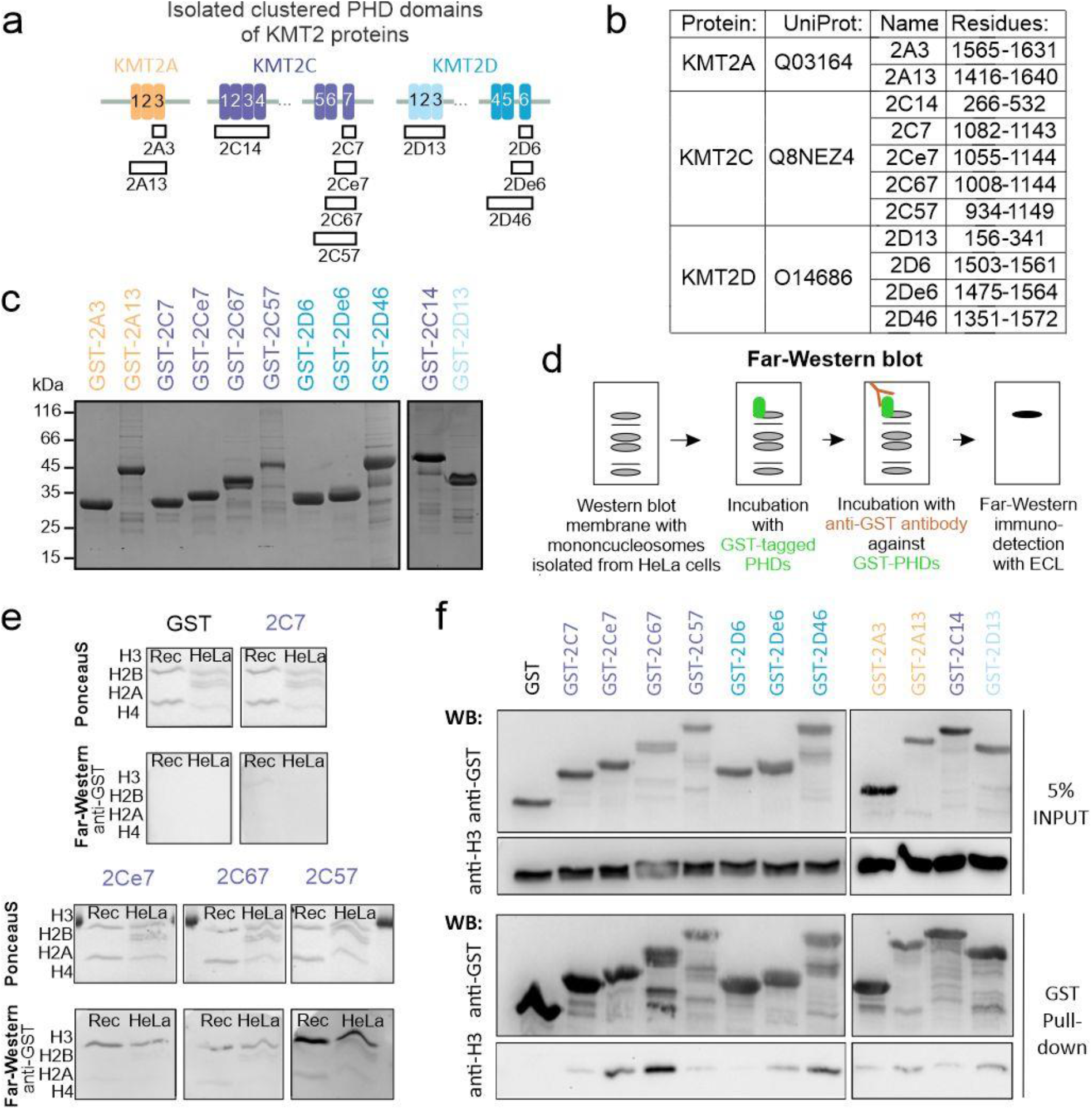
Affinity between PHDs and histones grows with the number of PHDs. **a** Overview of all purified GST-tagged PHDs. **b** Table with construct labels and protein regions, based on the UniProt entries. **c** 12% SDS-PAGE gel with 140 pmol purified isolated PHD domains stained with Coomasie blue. **d** Scheme of Far-Western blot methodology. **e** Far-Western blot performed with 100 nM solution of singlets (2C7, 2Ce7), doublet (2C67), and triplet (2C57) PHDs with membrane containing denatured recombinant H3/H4 (Rec) and endogenous histones isolated from HeLa S3 cells. All membranes were exposed simultaneously. **f** Western blot of the GST pull-down samples with singlets (2A3, 2C7, 2D6), enhanced singlets (2Ce7, 2De6), doublet (2C67), and full-length clusters (2A13, 2C14, 2C57, 2D13, 2D46) of GST-tagged PHDs. Bound mononucleosomes were identified using an anti-H3 antibody. The experiments were performed at the same time, at least with two replicates.

To characterise binding properties of PHD domains *in vitro* but within the nucleosome context, we used recombinant histones and endogenous mononucleosomes isolated from a HeLa S3 cell line (Suppl. Fig. 2a). First, we investigated the *in vitro* interactions with denatured proteins on the membrane by Far-Western blot (Fig. 2d-e, Suppl. Fig. 2b). The analysis indicated that all GST-PHDs, but not GST alone, bound to both recombinant and endogenous H3, whereas other histones were bound either weakly or not at all. The strength of the histone interactions grew with the number of PHDs (e.g., 2C7<2Ce7<2C67<2C57) (Fig. 2e). For the triplets and the quartet, the interaction was strongest for 2D46 and 2C57 and weaker for 2A13, 2C14, and 2D13 (Suppl. Fig. 2b). Histone modifications did not have a clear impact on the binding of PHDs, as both recombinant unmodified and endogenous modified H3 histones were bound with comparable affinity.

Next, we tested the interaction of PHDs with endogenous mononucleosomes in solution and performed GST pull-down assays (Fig. 2f). Consistent with the Far-Western results, we observed stronger interactions with PHD doublets and triplets than with singlets. In the Far-Western assay, the 2C57 triplet bound mononucleosomes better than the 2C67 doublet, but the opposite was observed in the pull-down assay. Possibly this discrepancy may be a consequence of lower stability of 2C57 during the longer overnight incubation used for the pull-down experiments. These results suggest that collaboration between the clustered PHD modules in KMT2A-D strengthens histone interactions.

### KMT2A-D clustered PHDs recognise specifically a subset of active promoters and enhancers

From our previous observations that full-length clustered PHD domains are stronger chromatin binders than their shorter fragments, we aimed to examine them in the cellular chromatin environment. Therefore, we generated stable HeLa S3 cell lines expressing NLS-EYFP-tagged triplet and quartet PHDs (NLS-EYFP-2A13, NLS-EYFP-2C14, NLS-EYFP-2C57, NLS-EYFP-2D13, and NLS-EYFP-2D46), as well as NLS-EYFP alone, as a negative control. In general, the recombinant proteins were weakly expressed (predominantly 2D46) (Fig. 3a, Suppl. Fig. 3a-d). Despite cell sorting after viral infection, the expression varied between cell lines and was anti-correlated with the strength of the interaction with mononulcoesomes in Far-Western and GST pull-down (Fig 2e, f), i.e., the strongest binder 2D46 was the least expressed, and 2D13 as one of the weakest binders was most highly expressed. We suspected that binding of the PHD domains to important chromatin regions affected the expression level and adjusted it to the tolerable levels. The EYFP fluorescence signal in the nucleus was diffuse, as for endogenous full-length KMT2D protein (Suppl. Fig. 3d), excluding protein overexpression artefacts from aggregation.

**Figure 3.**
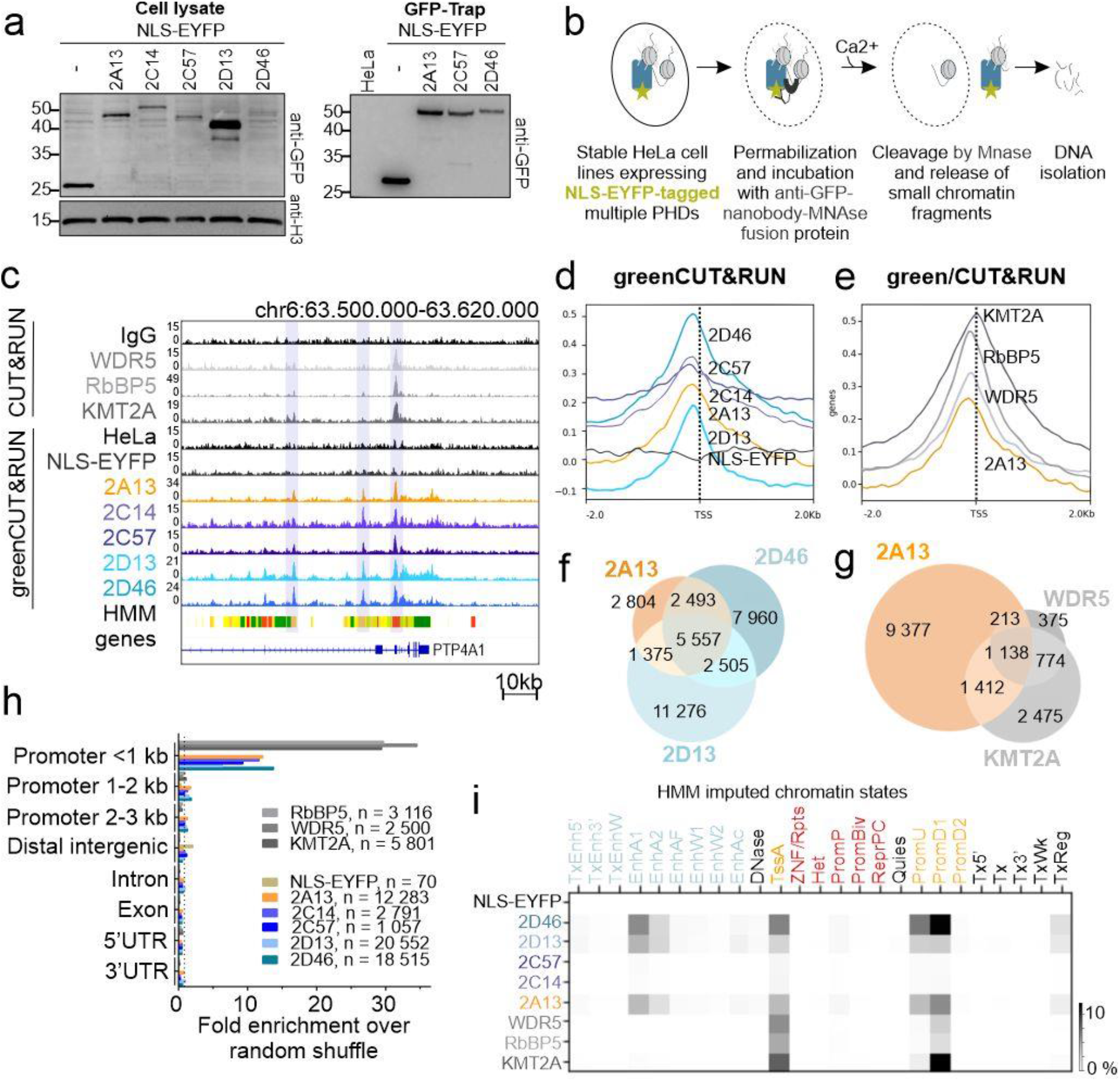
PHDs, expressed in the HeLa S3 cell lines, recognise specifically a subset of active promoters and to a lesser extent enhancers. **a** Western blot with anti-GFP antibody to detect NLS-EYFP-tagged clustered PHDs domains, expressed in stable HeLa S3 cell lines, generated by lentiviral transduction. Left panel - whole cellular lysate. Right panel - enriched proteins after GFP-Trap. **b** Scheme of greenCUT&RUN (gC&R) experiments. **c** IGV genome browser, showing read patterns of C&R hCOMPASS-like core subunits (WDR5, RbBP5 and KMT2A) and gC&R of clustered PHDs (2A13, 2C14, 2C57, 2D13, 2D46). The HMM lane shows imputed chromatin states; active promoter regions in red, active enhancers in yellow and orange and transcribed regions in green. **d-e** Heatmap plots comparing the distribution of reads around the TSS region (± 2 kb) of **d** clustered PHDs and NLS-EYFP (negative control) gC&R signals in comparison to the background of parental HeLa S3 cells and **e** hCOMPASS-like core subunits in comparison to IgG negative control. **f-g** Venn diagrams comparing MACS2 peaks of **f** selected PHDs (2A13, 2D13, 2D46) and **g** 2A13 with full-length hCOMPASS-like subunits (KMT2A, WDR5). Random shuffled data is shown on the Suppl. Fig. 3h. **h** Genomic distribution of PHDs and selected hCOMPASS-like subunits peaks. The quantification shows the enrichment in comparison to random shuffle. Distribution without enrichment is shown on the Suppl. Fig. 3j. **i** Jaccard analysis comparing the genome-wide distribution of peaks of selected proteins and HMM imputed chromatin states. Enhancers are in light-blue, promoters in yellow, poised or repressed chromatin states in red, and other chromatin states in black. The darker the signal, the stronger overlap is observed between the two samples. A detailed characterisation of imputed chromatin promoter and enhancer states with respect to chromatin marks is shown in Suppl. Fig. 5c.

In order to gain genome-wide insight into chromatin reader interaction patterns, we used generated stable cell lines for greenCUT&RUN (gC&R). gC&R relies on an MNase-anti-GFP nanobody, which cuts DNA around nucleosomes that are bound by an EYFP-tagged fusion protein of interest (Fig. 3b). For a better understanding of the role of PHD domains in KMT2A-D proteins recruitment, we also used standard C&R with antibodies against full-length KMT2 (KMT2A, KMT2C and KMT2D) proteins and against hCOMPASS-like subunits (WDR5, RbBP5). The antibodies against full-length KMT2C and KMT2D did not yield useful sequencing data. Surprisingly, read patterns of the gC&R data appeared similar across all clustered PHDs, when inspected in a genome browser (Fig. 3c). The NLS-EYFP negative control was comparable to the parental HeLa S3 background, indicating that the NLS-EYFP-tag did not cause artificial chromatin binding (Fig. 3c). Moreover, binding sites for PHDs and full-length proteins typically overlapped (Fig. 3c). Genomic heatmaps indicated strong enrichment of PHDs around transcriptional start sites (TSS), with the peak just upstream (~150-200nt) (Fig. 3d). We also observed a similar distribution for the investigated full-length hCOMPASS-like subunits. However, the sharp peak of KMT2A was shifted and located exactly at TSS (Fig. 3e). The nearly symmetrical peaks extended into the promoter region, but also downstream into the transcribed region.

We obtained good-quality sequencing data for NLS-EYFP-2A13 (~12 000 peaks), NLS-EYFP-2D13 (~20 500 peaks), NLS-EYFP-2D46 (~18 500 peaks) and moderate-quality sequencing data for NLS-EYFP-2C14 (~2 800 peaks) and NLS-EYFP-2C57 (~1 100 peaks). Importantly, a negative control of NLS-EYFP showed only around 70 peaks. In accordance with the observation from inspection in a genome browser, many of the PHDs binding sites were shared (~5 500 peaks for NLS-EYFP-2A13, NLS-EYFP-2D13, and NLS-EYFP-2D46) for the actual, but not randomly shuffled data (Fig. 3f, Suppl. Fig. 3e, f). For the full-length proteins we observed very well-defined, but fewer (2 500 - 5 800) binding sites than for the PHDs. These binding sites strongly overlapped with each other (~1 200 common peaks, Suppl. Fig. 3g) and partially with clustered PHDs (Fig. 3g, Suppl. Fig. 3h ~ 1 100 common peaks for NLS-EYFP-2A13, KMT2A, and WDR5, Suppl. Fig. 3i ~800 common peaks for NLS-EYFP-2A13, 2D13, 2D46, KMT2A, and WDR5). The overlap was absent upon random shuffle, confirming its relevance (Fig. 3g, Suppl. Fig. 3h).

Compared to randomly shuffled data, PHD domains and full-length proteins were 10 to 15-fold and 30 to 35-fold more enriched near promoters (<1kb), respectively (Fig. 3h, Suppl. Fig. 3j). To classify these binding sites more precisely, we assigned them to imputed chromatin states for HeLa S3 cells according to a “25 chromatin states” hidden Markov model (HMM) [34]. Overlap between genomic regions was quantified using the Jaccard index, defined as the ratio of the intersection and the union of two regions of interest. This analysis showed that all PHDs were bound at a subset of active promoters (active TSS (TssA), promoter downstream TSS 1 (PromD1), promoter upstream TSS (PromU), but not promoter downstream TSS 2 (PromD2)) and to lesser extent at a subset of active and strong enhancers (EnhA1, EnhA2) (Fig. 3i, Suppl. Fig. 3k). In contrast, the full-length subunits mapped exclusively to active promoters (mostly TssA and PromD1) (Fig. 3i). Taken together, these findings support the hypothesis that clustered PHDs play an important role in the targeting of the hCOMPASS-like complexes to active regulatory elements. Contrary to our initial expectation, all PHDs, also from enhancer-specific KMT2C-D proteins, are similarly enriched mostly at promoters and to a lesser extent at enhancers.

### Clustered PHDs bind H3K4me3 and H3 acetylation-rich loci, especially containing H3K9ac and H3K27ac, but not H4K16ac

To get a better insight into the characterization of loci bound by PHDs, we used a comprehensive set of HMM imputed chromatin marks [34]. First, we validated this set with experimental ChIP-Seq data from the ENCODE database [35], and our CUT&RUN data for H3K4me1 or H3K4me3 in HeLa S3 cells (Suppl. Fig. 4a, b). Next, we quantified the overlap between chromatin regions bound by PHD domains and HMM imputed histone marks, using the Jaccard overlap measure (Suppl. Fig. 4c, d). The analysis showed that clustered PHDs were specifically recruited to regions enriched with active promoter and enhancer marks (H3K4me1-3/ac, H3K9ac, and H3K27ac), but not to poised or silent chromatin marks, like H3K27me3 or H3K9me3 (Suppl. Fig. 4c, d). This pattern is consistent with bound imputed chromatin states (Fig. 3i). We also observed a strong correlation of PHD domain binding sites with other acetylation marks, like H2BK15ac, H3K23ac, and to a lesser extent H4 acetylation (H4K5ac, H4K8ac, H4K12ac) (Suppl. Fig. 4c, d). No such preference was detected for the NLS-EYFP negative control.

To confirm the strongest correlations of PHD binding sites determined by gC&R with imputed chromatin states, we selected antibodies against H3K4me1/3, H3K9ac, H3K23ac, H3K27ac, H3K56ac and H2BK15ac for the standard CUT&RUN. We also included H4K16ac, which was previously reported as a 2D6 target [25], as well as antibodies raised against multiple acetylated H2A.Zac and H4 histone peptides (H4K5acK8acK12ac (H4 3xKac) and H4K5acK8acK12acK16ac (H4 4xKac)). We discovered most acetylation marks were broadly distributed within actively transcribed gene bodies (Fig. 4a) with exceptions of H3K9ac and H3K27ac that were much more specific. Genomic heatmap plots indicated that H3 and H2B acetylation marks, and H3K4me3 signals peaked around the TSS, whereas H4 and H2A.Z acetylation signals had a noticeable dip in this position (Fig. 4b). Among all tested chromatin marks, H3K4me3, H3K9ac and H3K27ac were distributed most like PHD domains around the TSS (Fig. 3d, Fig. 4b). In addition, these three marks co-localized strongly with each other (Fig. 4c, Suppl. Fig. 4e, f), with PHD domain binding sites (Fig. 4d, Suppl. Fig. 5a, b) and chromatin states bound by the PHD domains (Fig. 3i, Suppl. Fig. 5c). In contrast, H4 acetylation marks, and particularly H4K16ac, co-localized poorly with PHD domains (Fig. 4d). Together, these observations are consistent with a role of H3K4me3, H3K9ac or H3K27ac, but not H4K16ac, as targets of clustered PHD domains in a cellular chromatin context.

**Figure 4.**
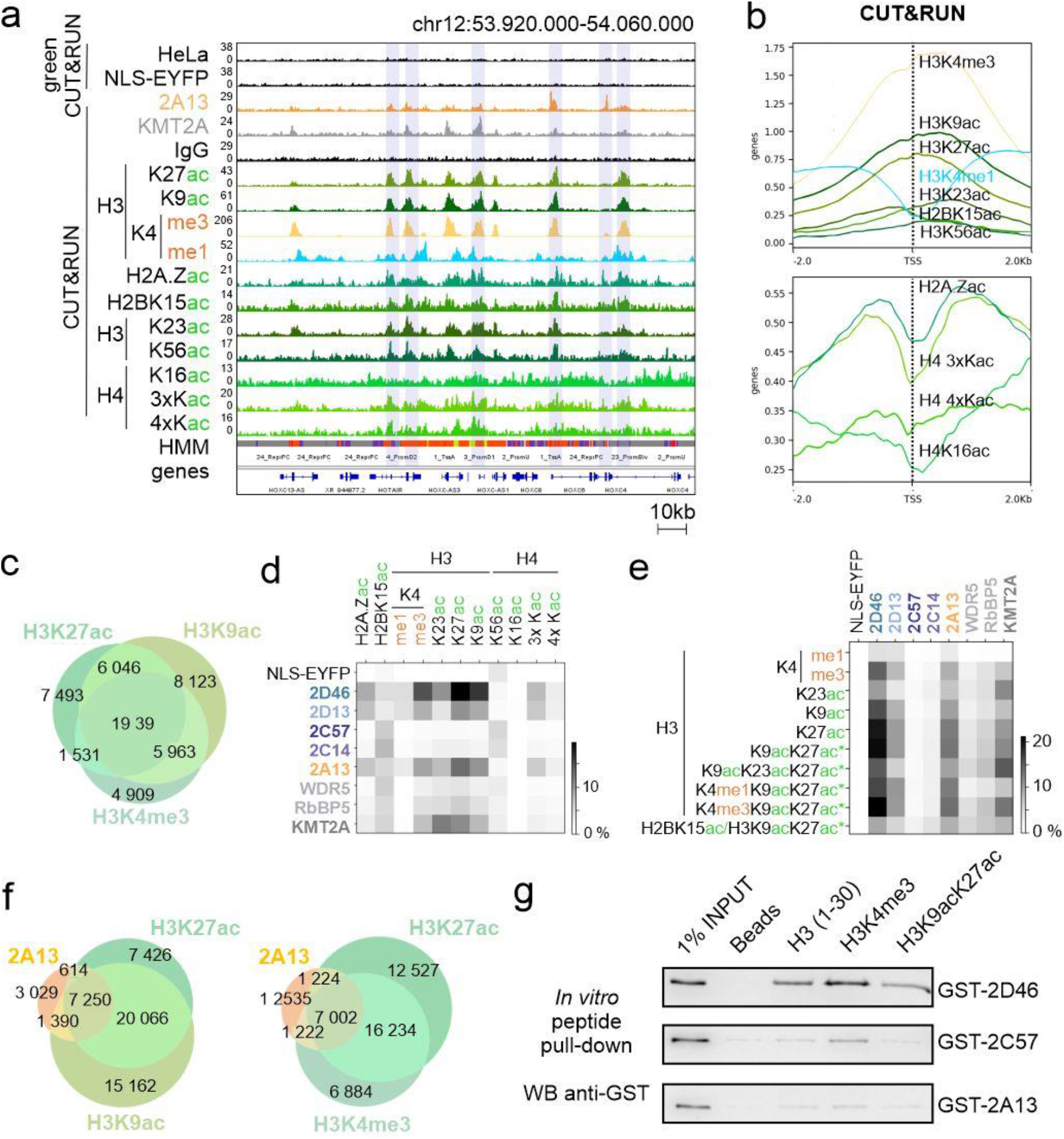
PHDs recognise loci enriched with multiple acetylation at H3 and H3K4me3, but not H4K16ac. **a** IGV genome browser snapshots of the reads distribution within the HOXC cluster of 2A13, NLS-EYFP gC&R, full-length KMT2A and selected histone marks (based on HMM imputed chromatin marks, Suppl. Fig. 4c): H3K4me1/3, H3K9ac, H2K23ac, H3K27ac, H3K56ac, H2A.Zac, H2BK15ac, H4K16ac, H4K16ac, H43xKac (H4K5acK8acK12ac) and H44xKac (H4K5acK8acK12acK16ac) C&R data. **b** Heatmap plots around TSS region (± 2 kb) of analysed histone marks in comparison to IgG negative control. Upper panel - H2B and H3 modifications. Bottom panel – H2A.Z and H4 modifications. **c** Venn diagram comparing peaks of the H3K4me3, H3K9ac and H3K27ac. **d** Jaccard analysis comparing the genome-wide peak distribution of PHDs or hCOMPASS-like subunits and of selected histone marks as well as **e** intersections (labelled with asterisk “*”) of the strongest correlated marks (H3K4me1/3, H3K9ac, H3K23ac, H3K27ac). **f** Venn diagrams comparing the distribution of peaks. Left panel - 2A13 PHDs,H3K9ac and H3K27ac. Right panel - 2A13, H3K27ac and H3K4me3. **g** Western blot with anti-GST antibody after biotinylated peptide *in vitro* pull-down with GST-tagged PHDs expressed in *E. coli*. Beads without peptide served as a negative control.

Overall, PHD domains, full-length KMT2A and hCOMPASS-like subunits (WDR5 and RbBP5) co-localized with similar chromatin marks – H3K4me3, H3K9ac, H3K27ac and H3K23ac. However, H3K23ac overlapped better with full-length proteins than with PHD domains (Fig. 4d). Additionally, we investigated the intersections of the strongest binders (H3K4me1/3, H3K9ac, H3K23ac and H3K27ac). We observed that the correlation of PHDs and full-length proteins with the combination of H3K4me3/H3K9ac/H3K27ac and H3K9ac/K23ac/K27ac, respectively, was stronger than with single modifications (Fig. 4e-f, Suppl. Fig. 5a). Taken together, the data suggests that the clustered PHD domains recognize multiple histone modifications. Biochemically, this is plausible, since each PHD domain may have separate methyl- and/or acetyl binding sites.

To probe a possible role of H3K4me3, H3K9ac or K3K27ac in the recruitment of PHDs *in vitro*, we examined interactions of the purified GST-tagged proteins (Fig. 2a) with synthetic histone tail peptides (H3 unmodified, H3K4me3, H3K9acK27ac) in a pull-down experiment (Fig. 4g). We observed that the PHD domains bound better to the H3K4me3 histone tail peptide than to the unmodified or the H3K9acK27ac doubly acetylated tail peptide (Fig. 4g). The experiment supports a direct role of H3K4me3 in PHD domain recruitment. It also suggests that the strong association of PHD domains with acetylated chromatin, seen in the gC&R data (Fig. 4d), could stem (in part) from indirect effects, such as increased histone tail peptide availability as a consequence of acetylation [36].

### A tandem of 2A13 PHDs with CXXC strongly improves promoter-specific recruitment

Surprisingly, the specificity of clustered PHDs from promoter-specific KMT2A and enhancer-specific KMT2C-D was similar (Fig. 3i). In full-length KMT2A, but not KMT2C-D, the clustered PHD domains are adjacent to other chromatin binding domains like the CXXC domain (binds CpG-rich DNA) or the bromodomain (binds acetylated histones). To probe domain cooperation and the effect on specificity, we aimed to generate stable HeLa cell lines featuring combinations of known chromatin binding domains of the KMT2A-D proteins. Unfortunately, we did not achieve a stable cell line with a construct covering all domains, possibly due to their strong interactions with chromatin and resulting toxicity. Instead, we obtained stable cell lines with very modest expression of artificial fusions of two clustered PHD domains from KMT2C (2C2P) or KMT2D (2D2P), a tandem of CXXC-PHD (2ACP) and cell lines with stronger expression of the KMT2C-D SET domains (2CS, 2DS) (Fig. 5a-b, Suppl. Fig. 6a-d). gC&R data for 2C2P, 2D2P, 2CS and 2DS did not indicate specific binding (Suppl. Fig. 6e), possibly due to a wrong linker between clustered PHDs in 2C2P and 2D2P construct.

**Figure 5.**
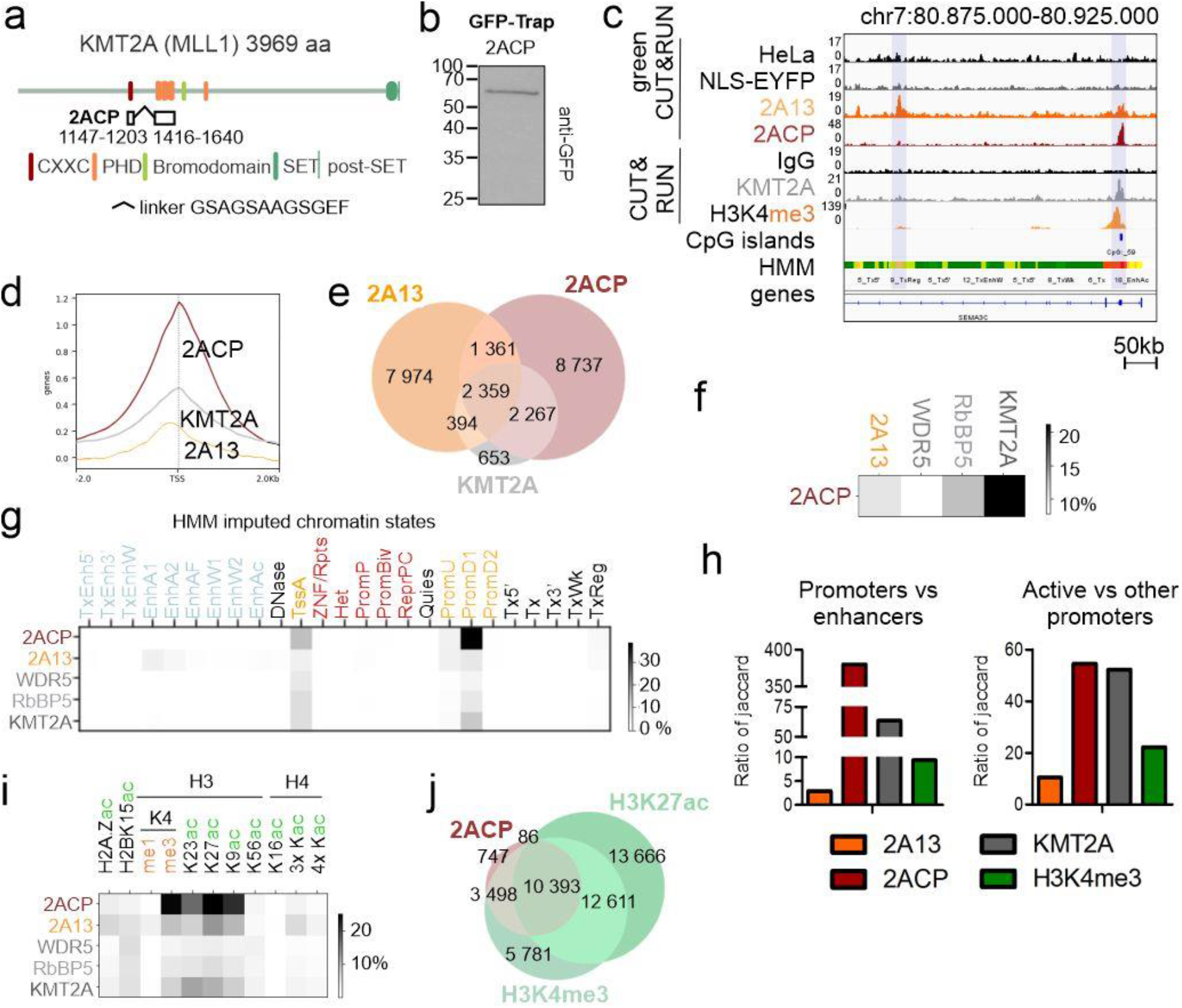
Tandem CXXC-2A13 of KMT2A protein improves promoter-specificity and overlap with full-length KMT2A in comparison to PHDs alone. **a** Scheme of the tandem CXXC-PHDs (2ACP) protein construct used for the generation of NLS-EYFP-2ACP stable HeLa S3 cell line by lentiviral transduction. **b** Western blot with anti-GFP antibody after GFP-Trap on NLS-EYFP-2ACP HeLa S3 stable cell lines. **c** IGV genome browser showing the distribution of reads within a representative gene (SEMA3C) generated by gC&R (2A13, 2ACP and negative controls HeLa, NLS-EYFP), CUT&RUN (KMT2A, H3K4me3 and negative control IgG), CpG islands (UCSC) and HMM chromatin states. **d** Heatmap plots around TSS region (± 2 kb) and **e** Venn diagram comparing binding sites of full-length KMT2A, tandem 2ACP and clustered PHDs 2A13. **f** Jaccard analysis comparing the genome-wide peak distribution of tandem 2ACP, full-length KMT2A, hCOMPASS-like subunits (WDR5, RBbP5) and clustered PHDs 2A13. **g** Jaccard analysis comparing the genome-wide distribution of the peaks of 2ACP, 2A13, full-length KMT2A, RbBP5, WDR5 with HMM imputed chromatin states. **j** Quantification of Jaccard ratios between loci enriched with indicated proteins (clustered PHDs 2A13, tandem CXXC-PHDs 2ACP, full-length KMT2A and H3K4me3). Left panel – ratio of all promoters (active TSS - TssA, promoter downstream TSS - PromD1/2, promoter upstream TSS - PromU, poised promoters - PromP, bivalent promoters - PromBiv) and enhancers (active enhancer - EnhA1/2, active enhancer flank - EnhAF, weak enhancer - EnhW1/2, H3K27ac possible enhancer - EnhAc, transcribed and enhancer - TxEnh5’/3’, transcribed and weak enhancer - TxEnhW). Right panel – ratio of active promoters (TssA, PromD1, PromD2, PromU) and other promoters (PromP, PromBiv). **i** Jaccard analysis comparing the genome-wide distribution of the peaks of 2ACP, 2A13, full-length KMT2A, RbBP5, WDR5 and previously selected (Fig. 3a, d) chromatin marks. Darker signal indicates a more pronounced overlap. Enhancers are in light blue, promoters in yellow, poised or repressed states in red, and other regulatory elements in black. **j** Venn diagram comparing binding sites of tandem 2ACP and with regions enriched for the most correlated histone marks (H3K4me and H3K27ac).

The 2ACP construct, a fusion of the CXXC and 2A13 PHD domains, recapitulated the KMT2A binding pattern much better than the 2A13 PHD domains alone (Fig. 5c-f). PHDs alone tended to be most highly enriched upstream to TSS (~150-200 bp) (Fig. 3d), contrary to 2ACP and full-length KMT2A which were most highly enriched directly at the TSS (Fig. 5d). 2ACP also bound a CG-rich DNA motif, in accordance with the known properties of CXXC domain (Suppl. Fig. 6f). Compared to 2A13, 2ACP had a stronger preference for promoters over enhancers, even exceeding the preference of full-length KMT2A (Fig. 5g, h). For 2ACP, the most correlated chromatin marks were H3K4m3, H3K9ac, K23ac, K27ac as for 2A13 (Fig. 5i), but the overlap was stronger (Fig. 5j). Taken together, these data shows that the promoter-specificity of KMT2A is due to the interplay of the PHDs (active promoters and enhancers) and CXXC (active promoters) chromatin reader domains.

### PHDs bind chromatin regions involved in cancer-related pathways and are often mutated in cancer patients

With gC&R and C&R data, we performed KEGG and gene ontology (GO) analysis for the chromatin targets of NLS-EYFP-2A13, NLS-EYFP-2D13, NLS-EYFP-2D46 PHDs, and full-length KMT2A (Fig. 6a, Suppl. Fig. 7a, b). Due to the small number of identified binding sites, we excluded 2C14 and 2C57 from this analysis. As a background control, we used H3K4me3 peaks, which showed the general active transcription in HeLa S3 cells. KEGG pathway analysis of bound genomic regions yielded similar results for all PHDs and full-length KMT2A protein (Fig. 6a, Suppl. Fig. 7a). Pathways with the lowest false discovery rate (FDR), not observed in the H3K4me3 control, were related to oncogenesis (e.g., “transcriptional misregulation in cancer” and “pathways in cancer”), in accordance with the known role of KMT2A-D and particularly of KMT2C as a tumour suppressor [37]. We observed the most significant associations with alcoholism and systemic lupus erythematosus, as previously observed for the HOXB1 transcription factor [38]. Results of the GO analysis of biological processes showed that most targets of clustered PHDs, but not H3K4me3, were associated with regulation of the mitotic cell cycle (Suppl. Fig. 7b). In the case of 2D13, developmental processes were also prominent.

**Figure 6.**
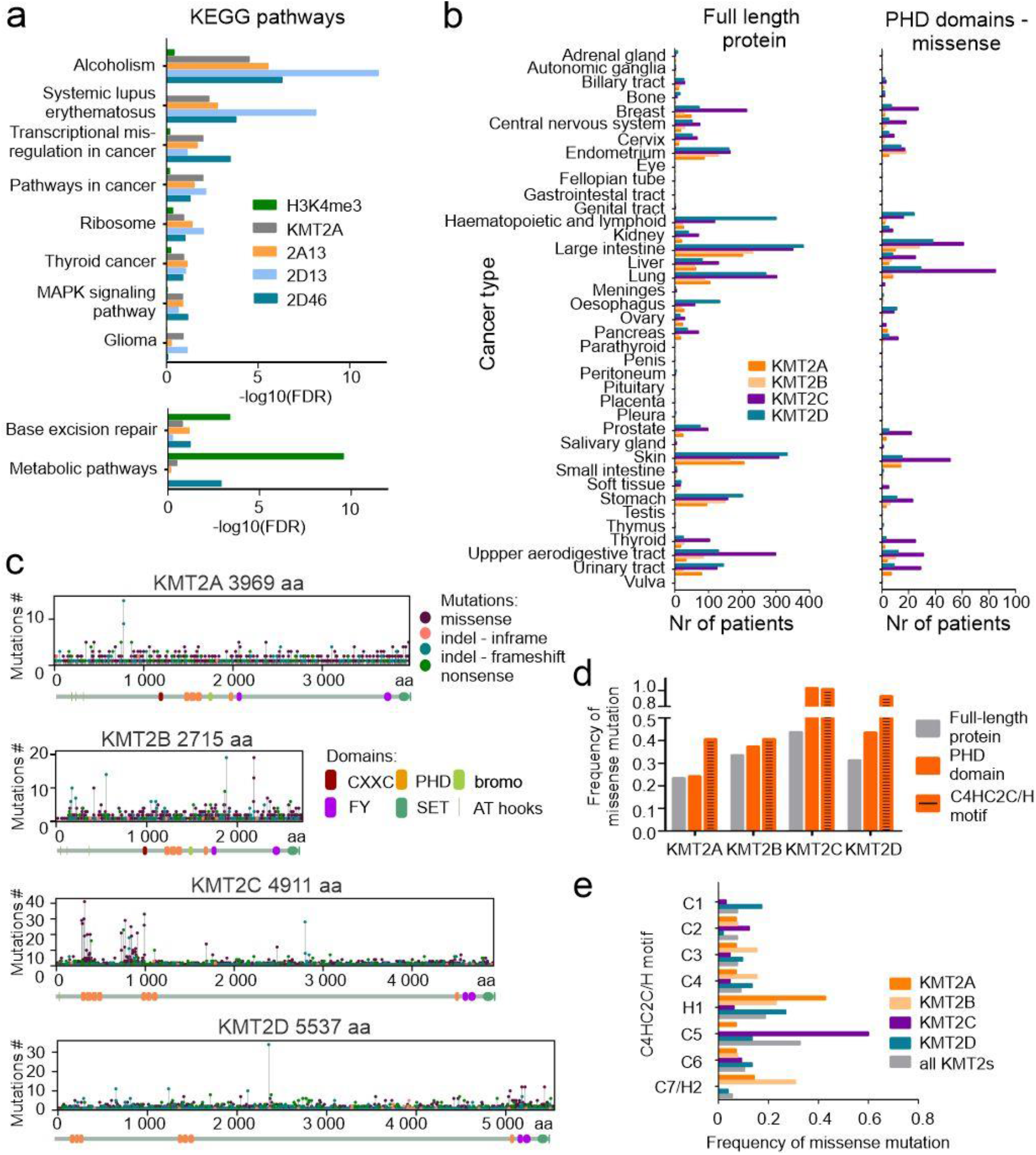
KMT2A-D PHDs bind loci involved in the cancer-related pathways. KMT2A-D PHDs, especially KMT2C, are mutated in cancer patients within crucial amino acids involved in the domain folding. **a** Top common hits of KEGG pathway of the peaks identified for full-length KMT2A (grey) and clustered PHDs. H3K4me3 serves as a control showing background of actively transcribed genes in HeLa S3 cells. FDR – false discovery rate. The top twelve hits for all proteins and GO biological processes are shown separately in Suppl. Fig. 7a, b. 2C14 and 2C57 were not included in the analysis due to the low number of identified binding sites. **b-e** Data obtained from the COSMIC database [70]. **b** Distribution of mutations among cancer patients. Left panel-missense, nonsense and in-del mutations within full-length KMT2A-D. Right panel – missense mutations within PHDs of KMT2A-D. Comparison of missense within full-length KMT2A-D and missense within PHDs is shown in the Suppl. Fig. 8. **c** Distribution of missense (violet), in-frame (pink), frameshift (light blue), and nonsense (green) mutations within KMT2A-D full-length proteins. **d** The ratio between missense mutations and the number of residues within a particular protein region (full-length, PHD, C4HC2C/H motif). Data is collected separately for KMT2A-D. **e** The frequency of missense mutations within the Zn^2+^ binding motifs (Cys4HisCys2Cys/His, C4HC2C/H motif) of the PHDs. Grey: mutations in all KMT2A-D proteins, orange: KMT2A, light orange: KMT2B, violet: KMT2C, blue: KMT2D.

As KMT2A-D proteins are frequently mutated in cancer [10], we used the COSMIC database [39] to gain more insight into the role of PHD domains. Within the database, among the KMT2A-D proteins, KMT2C is the most frequently mutated (~4 050 cases of cancer patients), followed by KMT2D (~2 600 cases), KMT2A (~1 580 cases), and finally KMT2B (~1 250 cases). In addition, KMT2A is most frequently affected by translocations that result in mixed lineage leukaemia (MLL). In contrast, point mutations in KMT2A-D proteins are most frequent in patients diagnosed with lung, liver, skin, or large intestine cancers (Fig. 6b, Suppl. Fig. 8). This pattern is similar when looking at mutations in full-length proteins or in PHD domains only.

Furthermore, our analysis of the COSMIC data for KMT2A-D showed that most of the mutations in cancer patients were found within the intrinsically-disordered regions (IDRs), which make up most of the proteins (Fig. 6c). The only exception to this trend was KMT2C. Mutations in this protein are enriched within the clustered PHDs, particularly the first cluster 2C14, as previously observed [10]. Some of the mutations in KMT2A-D alter crucial amino acids of the PHDs (especially within KMT2A and KMT2D proteins), specifically Cys or His residues (within the Cys4HisCys2Cys/His motif) that are needed for chelating Zn^2+^ ions and therefore domain folding (Fig. 6d-e). With an aim to see how these mutations within KMT2A and KMT2C-D PHD domains map to the protein surface, we generated AlphaFold-based models [40] and observed that the mutations tend to cluster spatially (Fig. 7a). Therefore, to verify the effect of most abundant cancer mutations within PHDs on their chromatin binding properties by gC&R, we chose 2A13 and 2D46 PHDs. We based our choice on the following criteria: the specific binding preferences of the wild-type domains (Fig. 3), and mutations within these domains are enriched in the C4HC2C/H motif, especially H1 (Fig. 6e). Thus, we generated mutated HeLa cell lines bearing single missense mutations (His1456Glu 2A13*, His1405Tyr 2D46*), localised in the folding motif (Fig. 7b-c). In all cases, peaks were drastically reduced compared to wild-type controls, confirming the expectation that the cancer mutations cause loss-of-function and reduce or abolish target specificity on chromatin (Fig. 7d).

**Figure 7.**
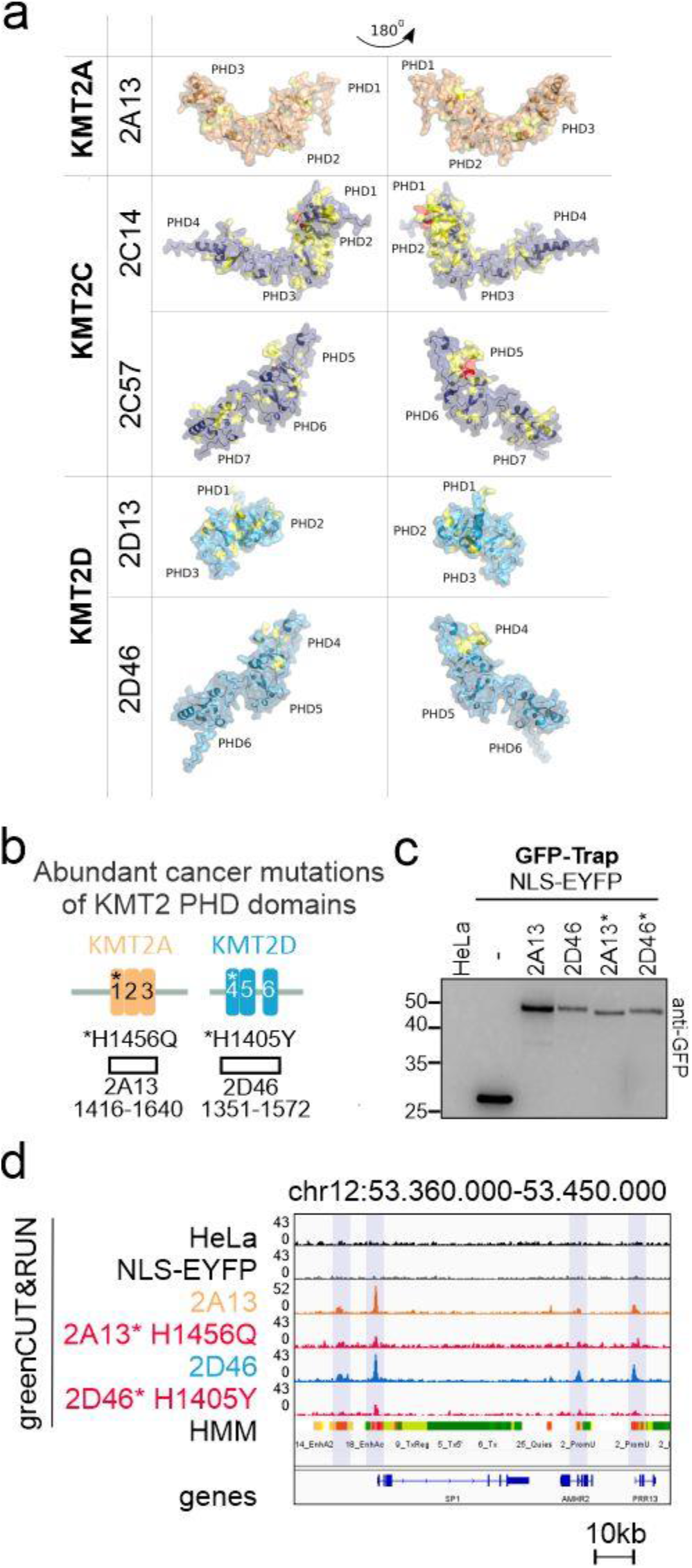
Mutations of PHD domains found in cancer patients impair chromatin reader properties. **a** Structure of PHDs predicted by Alphafold with COSMIC mutations mapped to the protein surface. Mutations found in 1-10 or more COSMIC patients are shown in yellow and red, respectively. **b** Scheme of the clustered PHDs of KMT2A-D proteins with mutations selected for experimental analysis, **c** Western blot with anti-GFP antibody after GFP-Trap of NLS-EYFP, 2A13, 2D46 and of their cancer mutation variants. **d** IGV genome browser snapshot illustrating impaired chromatin binding of the mutated PHD domains (2A13* H1456Q and 2D46* H1405Y) in comparison to the wild-type 2A13 and 2D46 domains.

## DISCUSSION

### GreenCUT&RUN as a tool to characterize chromatin readers

Chromatin reader domains are frequently studied *in vitro* using peptide pull-down and histone arrays with short and modified histone peptides [41–43]. These approaches are problematic, because they do not take into account the chromatin context. Recently, synthetic nucleosomes with defined chromatin modifications have become commercially available as an alternative to synthetic histone tail peptides for research on chromatin interactions [44]. Experiments with such nucleosomes have demonstrated that some histone tail modifications affect chromatin reader binding indirectly, by modulating histone tail availability [36,45]. The synthetic nucleosomes with defined posttranslational modifications have already been used for a so-called readerCUT&RUN assay [45]. Although this assay is more physiological than the assays with synthetic histone tail peptides, it still relies on reconstituted nucleosomes. ChIP-Seq and gC&R or C&R experiments are currently the preferred methods to characterize chromatin interactions in a physiological context. However, in our research only gC&R is applicable for the clustered PHDs. Compared to the more traditional ChIP-Seq experiments, both gC&R and C&R experiments require fewer reads and benefit from a much better signal to noise ratio. We therefore believe that gC&R and C&R experiments could become a useful tool to characterise chromatin readers.

### Clustered PHDs of KMT2A-D bind chromatin regions enriched in H3 acetylation and H3K4me3

In this study, we observed strong overlap of PHDs with multiple acetylation marks (Fig. 4d, Suppl. Fig. 4c, d). These acetylation marks are all expected to be found in active chromatin, in line with the biological role of epigenetic memory proteins, like KMT2A-D (Fig. 8a). Among genomic elements of the same type, PHDs appear to discriminate chromatin states based on their acetylation levels. Preferentially bound promoter PromD1 is highly acetylated while unbound promoter PromD2 is not (Suppl. Fig. 5c). Highly acetylated enhancer EnhA1 is preferred by PHDs over less acetylated enhancer EnhA2 (Suppl. Fig. 5c). We also observed that clustered PHDs peaked more sharply around the TSS than single chromatin marks (Fig. 3d ~1kb, Fig. 4b ~2-3 kb wide). This finding suggests that PHDs bind several chromatin marks simultaneously.

**Figure 8.**
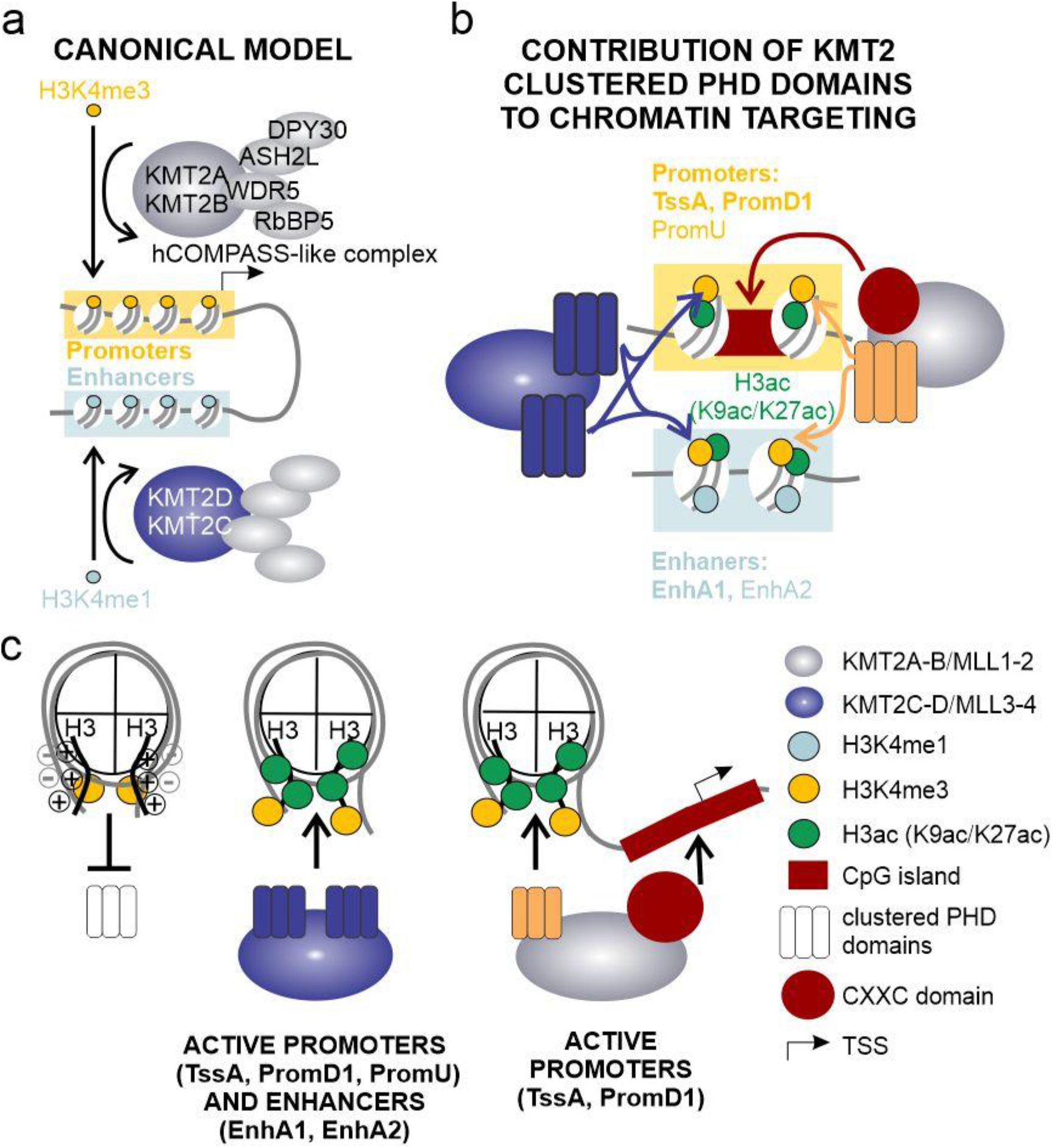
Involvement of PHDs in targeting of hCOMPASS-like complexes. **a** Canonically, KMT2A-B are involved in recognizing active promoters and catalyzing H3K4me3. In contrast, KMT2C-D are shown to target active enhancers and introduce H3K4me1. **b** We show that chromatin reader domains, such as clustered PHDs, are involved in targeting KMT2A-D proteins to active promoters (in particular around TSS (TssA) and within promoter PromD1) and to a lesser extent to active enhancers (EnhA1, EnhA2) rich in H3K4me3, H3K9ac and K3K27ac. The CXXC domain is present only within promoter-specific KMT2A and KMT2B, but not enhancer-specific KMT2C and KMT2D. It is highly attracted by non-modified CpG islands (any single modification repels this interaction), and enhances KMT2A-B promoter preference. **c** Scheme of nucleosome interactions between H3 histone tail, DNA and reading domains (PHDs and CXXC). Left panel – positively charged H3 tail interacts with dyad DNA, so that H3K4me3 is hidden and PHD domains are not bound. Middle panel – acetylated H3 tail (i.e., containing H3K9ac and H3K27ac) loses positive charge, so the H3K4me3 is available for the interaction with PHDs. Right panel – in the case of promoter-specific KMT2A we observe cooperation of two reading domains – PHDs reading available H3K4me3 and CXXC reading unmodified CpG island within nucleosome free region (NFR) at TSS.

Comprehensively, our gC&R and C&R data shows that H3K4me3, acetylation marks H3K9ac and H3K27ac and their intersections have the highest overlap score in the Jaccard analysis with PHDs (Fig. 4d-f). These marks are also distributed most similarly to PHDs around the TSS (Fig. 3d, Fig. 4b). Our data does not support the previously reported binding of singlet 2D6 to H4K16ac. This interaction was characterized using synthetic histone peptide [31] (Fig. 4a, b, d), that may not faithfully recapitulate the interactions with a full nucleosome.

H3K9ac and H3K27ac distinguish active from poised promoters and enhancers [46,47]. Enrichment of the PHDs at these sites is therefore consistent with a preference of the PHD domains for active promoters and enhancers (Fig. 3i, Fig. 8b). However, *in vitro* pull-down experiments indicate similar affinity of the PHD domains to unmodified and H3K9acH3K27ac histone peptide tails (Fig. 4g). It is possible that acetylation influences PHD domain binding indirectly in a chromatin context, as H3 tails are localised near the entry and exit sites for DNA dyad [48]. Hence their acetylation could strongly decrease interactions with nucleosomal and linker DNA, and thereby increase H3K4me3 availability (Fig. 8c), as has been proposed for the PHDs of BPTF [36].

PHD domains of promoter-specific KMT2A and enhancer-specific KMT2C and KMT2D differ considerably (conservation 40-50%, Suppl. Fig. 1b). Yet, all of them are recruited to well-defined H3K4me3 and H3 acetylation-rich active promoters and enhancers (Fig. 3i). The acetylation marks that overlap best with PHD domain binding sites, H3K9ac and H3K27ac, are known as both promoter and enhancer mark. By contrast, H3K4me3 is canonically associated with promoter regions. However, H3K4me3 was also shown to be present at strong enhancers [49]. In our data, H3K4me3 is found in ~23% and ~10% of the EnhA1 and EnhA2 active enhancer regions, respectively (Suppl. Fig. 9a). By contrast, H3K4me3 decorates 76% TssA, 90% PromD1, 70% PromD2 and 61% PromU (Supp. Fig. 9b). In general, 2A13 binds around 10% and 5% of EnhA1 and EnhA2, respectively (Suppl. Fig. 9c). However, among the H3K4me3 decorated EnhA1 and EnhA2 enhancers this overlap is several folds bigger, as 30% (1729 out of 5717) and 25% (791 out of 3135) are bound by 2A13, respectively (Suppl. Fig. 9b). Together, these data confirms the observation that clustered PHDs recognize both active promoters and enhancers by targeting regions marked by H3K4me3 and H3ac (especially H3K9ac and H3K27ac) (Fig. 8c).

### The CXXC domain strongly enhances the promoter preference of KMT2A

The surprisingly similar binding preferences of PHD domains from promoter- and enhancer specific KMT2A-D prompted us to focus on the overall architecture of these proteins. Interestingly, only promoter-specific KMT2A-B, but not enhancer-specific KMT2C-D, contain bromodomains and CXXC domains (Fig. 1a). Bromodomains typically bind acetylated lysine residues [50], associated with a permissive chromatin state [51]. They could therefore potentially help to target methyltransferases to active chromatin, thus facilitating positive transcriptional memory. Currently, there is no evidence that the bromodomains could contribute to the preference of KMT2A-B for promoters over enhancers. By contrast, KMT2A CXXC domains bind only at unmodified CpGs and are repelled by any single cytosine modification [12,23]. Unmodified CpGs are enriched in CpG islands (CGIs) mostly within active promoters, but not elsewhere in the genome [24].

The gC&R data in this study shows that an artificially fused tandem of CXXC and 2A13 strongly recapitulates the promoter-specific properties of full-length KMT2A (Fig. 5a-h). We also observed that 2A13 PHDs peak upstream of the TSS (~150-200nt), whereas 2ACP and full-length KMT2A peak directly at the TSS. We attribute the TSS upstream binding of 2A13 (and other PHDs) to the nucleosome free region (NFR) directly at the TSS of the transcribed genes [52]. In contrast, 2ACP and full-length KMT2A can bind directly at the TSS because the CXXC domain binds CG-rich DNA directly and is not dependent on the presence of nucleosomes (Fig. 8c right panel, Suppl. Fig. 6f). Additionally, the gC&R data indicates significantly higher promoter to enhancer preference for 2ACP than for 2A13 or even full-length KMT2A (Fig. 5g, h). Therefore, our findings strongly suggest that the presence of the CXXC domain in KMT2A-B, but not KMT2C-D accounts for the observed stronger promoter preference of KMT2A-B compared to KMT2C-D (Fig. 8c). As KMT2C-D catalyse H3K4me1 [53,54], present in both enhancers and promoters, we believe that their PHD domains may indeed target both chromatin states (active enhancers and promoters), not exclusively just the active enhancers (Fig. 8).

### Chromatin targeting by other hCOMPASS-like subunits

KMT2A-D proteins are recruited to their chromatin targets not only by their reading domains but their localization can also be influenced by their association with other chromatin reader subunits of hCOMPASS-like complexes. For WDR5, a preference for methylated H3K4, particularly dimethylated H3K4 [14,55,56] with modulation by methylation of H3R2 [19] and serotonylation of H3Q5, was demonstrated biochemically [57]. For ASH2L, strong evidence for DNA binding was shown [18,58]. Menin, a component of hCOMPASS-like complexes with KMT2A-B catalytic core, was observed to bind a plethora of transcription factors [59]. Despite all these influences on KMT2A-D targeting, our data shows that the 2A13 triple PHD domain of KMT2A and the CXXC-2A13 tandem bind to similar targets as full-length KMT2A in the hCOMPASS-like complex context, supposedly making an important contribution to general protein targeting.

### Clustered PHDs in KMT2A-D proteins are conserved and developmentally required in animals

The clustered PHD domains are among the most highly conserved regions of KMT2A-D proteins (Fig. 1b). They occur in vertebrate and invertebrate KMT2A-D otrthologues. The *Drosophila melanogaster* orthologue of KMT2A-B, Trx, has one PHD triplet. The *Drosophila* orthologue of KMT2C-D, Trr, lacks clustered PHDs but interacts with a separate protein known as lost PHDs (*Lpt* or *Cmi*) that contains two clusters of PHDs (a quartet and a triplet). In Drosophila, ablation of *Cmi* results in larval lethality [60], at least in part due to defects in pattern formation [61]. This observation highlights the crucial role of clustered PHDs in organism development. To our knowledge, triplets and quartets of PHD domains are unique to the animal KMT2A-D proteins and not found elsewhere.

### The clustered PHDs in KMT2A-D proteins are frequently mutated in cancer

Missense and nonsense mutations, as well as small deletions and insertions, are frequent in KMT2A-D and some of them map to the PHDs (Fig. 6b, c, Fig. 7a). Enrichment is seen with strong statistical significance for the frequently mutated KMT2C, as observed previously [11]. The distribution of mutations to many sites, rather than a few selected sites, and the enrichment of alterations in conserved residues of the PHDs suggest that the mutations cause a loss-of-function and change-of-function, respectively [62]. The accumulation of alterations in conserved residues of the PHDs suggest that PHD dysfunction is an important contributor in KMT2A-D driven malignancies. This hypothesis is supported by our observation of frequent mutations within crucial amino acids for the folding of the PHD domains (Fig. 6d-e) and by the demonstration that the mutations disrupt domain targeting (Fig. 7b). However, KMT2A alterations in leukaemia, particularly mixed lineage leukaemia, are not small local changes, but chromosomal translocations generating novel fusion proteins. Importantly, translocation breakpoints cluster into a major Breakpoint Cluster Region (BCR) with most breakpoints in the core region from exon 9 to 11 [63,64], and a minor breakpoint cluster region from exons 19 to 26 [65]. As the coding region for clustered PHDs of KMT2A (2A13) starts in exon 11, 2A13 is either completely absent or only present in a truncated form within the oncogenic fusion proteins. Thus, we conclude that biology of PHDs is relevant for KMT2A-D loss-of-function malignancies, but not for translocation-driven KMT2A gain-of-function leukaemias.

In conclusion, we have shown that clustered PHD domains direct KMT2A and KMT2C-D to a subset of active promoters (in particular in close proximity to TSS) and to a lesser extent to active enhancers of cancer-related pathways (Fig. 8b, c). PHDs bind the chromatin loci in the multiple binding mode by recognizing regions enriched notably with H3K4me3 and H3ac (H3K9ac and H3K27ac). In our experiments, we have shown that mutations in the PHD domains found in cancer patients disrupt targeting of the isolated domains. It is likely that these mutations are selected in cancer because they also disrupt or at least negatively impact targeting of the entire hCOMPASS-like complexes. Hence, our experiments suggest a role of the PHD domains of KMT2A-D proteins in the maintenance of positive epigenetic memory.

## MATERIALS AND METHODS

### Cloning and overexpression of PHD domains

*E. coli* codon-optimised synthetic genes (GeneArt, ThermoScientific; Table S1) encoding the PHD domains of human KMT2A (UniProt: Q03164, 2A3, aa 1565–1631; 2A13, aa 1416–1640), KMT2B (UniProt: Q9UMN6, 2B13, aa 1182–1414), KMT2C (UniProt: Q8NEZ4, 2C14, aa 266–532; 2C7, aa 1083–1143; 2Ce7, aa 1055–1144; 2C67, aa 1008–1144; 2C57, aa 934–1149) and KMT2D (UniProt: O14686, 2D13, aa 156–341; 2D6, aa 1503–1561; 2De6, aa 1475–1564; 2D46, aa 1351–1572) were cloned using BamHI and XhoI restriction sites into pGEX-P6-2 plasmids encoding a N-terminal GST-tag (Fig. 2b). For GST overexpression, an empty pGEX-P6-2 plasmid was used. In addition, human codon-optimised synthetic genes (GeneArt, ThermoScientific; Table S1) encoding the clustered PHD domains of KMT2A (UniProt: Q03164, 2A13, aa 1565–1631), KMT2C (UniProt: Q8NEZ4, 2C14, aa 266–532; 2C57, aa 934–1149) and KMT2D (UniProt: O14686, 2D13, aa 156–341; 2D46, aa 1351–1572) were cloned using the SLIC method [66,67], first to pEYFP plasmids and then into a customised pLJM1-EGF1-based plasmid encoding N-terminal EYFP for the generation of a stable cell line by lentiviral transduction. Briefly, NLS-EYFP-PHD fusion gene fragments were amplified with the primers specified in Table S2 using Q5 DNA Polymerase. Plasmid pLJM1 was linearised using AgeI and EcoRI restriction enzymes. DNA fragments were incubated for 30 minutes with T4 DNA polymerase at 22 °C to generate overhangs. The reaction was stopped by the addition of dCTP to a 2 mM final concentration. Inserts and vector were mixed in a 1:1 molar ratio and incubated for 30 minutes at 37 °C before being transformed into NEB stable *E. coli* competent cells.

Other human codon-optimised synthetic genes (GeneArt, ThermoScientific; Table S1) encoding tandem reading domains KMT2A (UniProt: Q03164, 2ACP, aa 1147-1203 and 1416-1640), KMT2C (UniProt: Q8NEZ4, 2C2P, aa 266-532 and 934-1149), KMT2D (UniProt: O14686, 2D2P, aa 156-341 and 1351-1572) fused with the artificial linker GSAGSAAGSGEF, SET domains of KMT2C (UniProt: Q8NEZ4, 2DS, aa 4751-4911) and KMT2D (UniProt: O14686, 2DS, aa 5371-55537) and cancer-related mutations in KMT2A PHDs (UniProt: Q03164, 2A13* H1456Q, aa 1416-1640), KMT2D (UniProt: O14686, 2D46* H1405Y, aa 1351-1572) were cloned using using BamHI and XhoI into a customised pLJM1-EGF1-based plasmid encoding N-terminal EYFP for the generation of a stable cell line by lentiviral transduction.

All protein overexpression was performed in *E. coli* BL21(DE3) pLys cells. The cells were grown in LB media supplemented with 100 ng/ml ampicillin at 37 °C until OD_600_ reached 0.6/cm–0.8/cm. Cells were then transferred to 4 °C for 30 minutes. After induction with 1 mM IPTG and in the presence of 150 μM ZnSO_4_, proteins were expressed overnight at 22 °C. The cells were then harvested and stored at −80 °C in pellets from 1L of culture batches.

### Protein purification

Frozen cells overexpressing GST or GST-PHDs were resuspended in 40 ml ice-cold lysis buffer (50 mM Tris-HCl pH 7.5, 400 mM NaCl, 5% glycerol, 150 μM ZnSO_4_, 1 mM DTT) supplemented with 0.25 mM PMSF. After sonication, the lysate was cleared by ultracentrifugation for 30 minutes. Next, the supernatant was applied to Glutathione Sepharose™ 4B beads (GE Healthcare) equilibrated with lysis buffer. After 2 hours incubation, the beads were extensively washed with washing buffer (50 mM Tris-HCl pH 8.0, 400 mM NaCl, 5% glycerol, 150 μM ZnSO_4_, 1 mM DTT). Protein was eluted by incubating the beads with elution buffer (200 mM Tris-HCl pH 8.0, 150 mM NaCl, 5% glycerol, 20 mM reduced L-glutathione, 150 μM ZnSO_4_, 1 mM DTT) for 15 minutes. The purified protein was concentrated using Amicon Ultra centrifugal filters with 10 kDa cut-off (Millipore) and dialyzed overnight against storage buffer (50 mM Tris pH 8.0, 150 mM NaCl, 150 μM ZnSO_4_, 1 mM DTT, 50% glycerol). Protein concentrations were determined spectrophotometrically (Nanodrop, Thermo Fisher Scientific). Aliquoted samples were frozen in liquid nitrogen and stored at −80 °C.

### Isolation of mononucleosomes

Mononucleosomes were isolated from HeLa S3 cells. A pellet of 20 million HeLa S3 cells was resuspended in 5 ml of ice-cold TM2 buffer (10 mM Tris pH 7.5, 2 mM MgCl_2_, 0.5 mM PMSF). Next, NP-40 was added to a final concentration of 0.6%. After incubation on ice for 5 minutes, the pellet was washed with TM2 buffer, resuspended in 400 μl of TM2 buffer containing micrococcal nuclease (New England Biolabs) and 1mM CaCl_2_, and incubated in a Thermomixer for 10 minutes at 37 °C. Subsequently, 2 mM EGTA, 300 mM NaCl, and 0.1% Triton X-100 were added, and the mixture was transferred into Bioruptor tubes (Diagenode). After sonication for 5 cycles of 30 seconds ON/OFF at 4 °C (Bioruptor Pico, Diagenode), the mixture was centrifuged (13 000xg) for 10 minutes at 4 °C, and the supernatant was transferred to a fresh tube. For quality control, DNA was isolated from a small aliquot (PCR clean up, A&A Biotechnology) and checked on 1.5% agarose gel for DNA length (~150 nt). Finally, mononucleosomes were divided into aliquots, fast-frozen with liquid nitrogen, and stored at −80 °C.

### Human cell lines

HeLa S3 cells, HeLa S3-derived stable cell lines, and HEK293T cells were cultured in Dulbecco’s Modified Eagle Medium (DMEM, Biowest) supplemented with 10% foetal bovine serum (FBS, Biowest) and penicillin-streptomycin solution (Biowest) at 37 °C in an humidified 5% CO_2_ atmosphere. Cells were passaged using trypsin and stored in 5% DMSO and DMEM containing 10% FBS and antibiotics at −80 °C.

### Generation of stable cell lines

Stable HeLa S3 cell lines expressing NLS-EYFP-tagged proteins were generated using lentiviral transduction. Lentivirus stocks were prepared using plasmids pMD2.G and psPAX2, and a pLJM1-based transfer plasmid, by standard transfection with PEI (Sigma Aldrich #408727). HEK293T cells were grown in a 15 cm dish following transfection. Upon reaching ~90% cell confluency, the medium was aspirated and replaced with fresh medium without antibiotics. Transfer plasmid pLJM1 and plasmids psPAX2 and pMD2.G were mixed in 4:2:1 ratio in Opti-MEM (20 μg, final volume 0.7 ml). Next, PEI in Opti-MEM was added (60 μg in a total volume 0.7 ml). The transfection mixture was briefly vortexed, incubated for 20 minutes at room temperature, and slowly added to the cell culture. 17 hours post-transfection, the medium was exchanged for fresh DMEM supplemented with 25 mM HEPES, 4 mM caffeine, and 1 mM sodium butyrate. The medium was harvested 48 hours post-transfection and centrifuged for 3 minutes at 3 000 g. The supernatant was then filtered through a 0.45 μm PES filter, aliquoted, and stored at −80 °C.

To generate stable cell lines, 0.1 million HeLa S3 cells were seeded per well in a 6 well plate 24 hours before infection. Infection was carried out for 72 hours. At every 24 hours, media was exchanged for a new virus aliquot and fresh media in a ratio of 2.5:0.5, supplemented with DEAE-dextran (6 μg/ml). The negative control NLS-EYFP HeLa S3 stable cell line was infected once after 24 hours. 72 hours after the first virus addition, the cells were transferred to a 10 cm plate and maintained in media supplemented with 2 μg/ml puromycin. Non-infected parental HeLa S3 cells were also treated with puromycin to confirm successful selection.

### Flow cytometry and cell sorting

After successful puromycin selection, stable cell lines were sorted using a BD FACSAriaII (Becton Dickinson), in collaboration with Microscopy and Cytometry Facility in IIMCB. Briefly, parental and stable HeLa S3 cell lines were washed with PBS, harvested with trypsin-EDTA, washed twice with PBS, resuspended in PBS buffer and transfered to 5 ml FACS tubes. Next, cells expressing EYFP above background levels of parental HeLa S3 cell line were sorted out and transferred back to the culture plate. Sorted cells were cultured in standard conditions for at least 2 weeks to reach confluency, re-analyzed using BD FACSAria II cell sorter and used for the gC&R experiments. To detect EYFP fluorescence signal in the cells blue laser 488 nm and FITC emission filter (530/30) were used. Finally, the cells were stored in 5% DMSO and DMEM containing 10% FBS and antibiotics at −80 °C. Flow cytometry data was plotted and analyzed using FlowJo software (Becton Dickinson).

### GST pull-down, peptide pull-down and GFP-Trap

For GST pull-down, purified GST-tagged PHD domains (1 μM) were incubated with 30 μg of isolated mononucleosomes in 500 μl of pull-down buffer (10 mM Tris pH 7.5, 150 mM NaCl, 0.5% Triton X-100, 1 mM DTT, 150 μM ZnSO_4_, 1 mM PMSF, 1xEDTA-free protease inhibitors [Roche]) overnight on a rotating wheel at 4 °C. 10% of the mixtures were put aside as input. Next, 40 μl of Glutathione Sepharose™ 4B beads (GE Healthcare) were washed three times with pull-down buffer and added to the sample containing the PHD mononucleosome complexes. The mixtures were then incubated for 2 hours on a rotating wheel at 4 °C. Following this, the supernatants were collected as flow through and the beads were washed three times with washing buffer (10 mM Tris pH 7.5, 150 mM NaCl, 0.5% Triton X-100). Finally, the beads were incubated at 95 °C with SDS-loading buffer (50 mM Tris pH 6.8, 2% SDS, 5% β-mercaptoethanol, 10% glycerol, bromophenol blue) for 10 minutes for Western blot.

For peptide pull-down, first 25 μl Dynabeads™ MyOne™ Streptavidin C1 were incubated with 250 pmols of selected biotinylated peptides (GeneScript, Supplementary Table X) in 100 μl of peptide binding buffer (50 mM Tris pH 8.0, 300 mM NaCl, 0.1% NP-40, 1mM PMSF) for 30 minutes on a rotating wheel at room temperature. Next, resins were washed three times and incubated with GST-tagged PHD domains (25 pmol) in 300 μl protein binding buffer (50 mM Tris pH 8.0, 150 mM NaCl, 0.1% NP-40, 0.5% BSA, 1 mM DTT, 150 μM ZnSO4, 1 mM PMSF) for 4 hours on a rotating wheel at 4 °C. 1% of the mixtures were put aside as input. Following this, the beads were washed three times with the protein binding buffer. Finally, the beads were boiled at 95 °C with SDS-loading buffer (50 mM Tris pH 6.8, 2% SDS, 5% β-mercaptoethanol, 10% glycerol, bromophenol blue) for 10 minutes for Western blot.

For GFP-Trap, 10 cm plates were seeded with the selected NLS-EYFP-tagged stable HeLa S3 cell line. When the plate reached the full confluency (~10^7^ cells), the cells were washed with ice-cold PBS and scraped from the plate. Next, cells were washed two more times with ice-cold PBS and lysed with RIPA buffer lacking EDTA (10 mM Tris 7.5, 150 mM NaCl, 0.1% SDS, 1% Triton X-100, 1% deoxycholate), supplemented with DNaseI, 2.5 mM MgCl2, 1mM PMSF and EDTA-free protease inhibitor cocktail (Roche) for 30 minutes on ice. The lysate was centrifuged at 20 000xg for 10 minutes at 4 °C, diluted with the dilution buffer (10 mM Tris pH 7.5, 150 mM NaCl, 1mM PMSF) and mixed with 10 μl equilibrated GFT-Trap magnetic agarose beads (Chromotek). The lysate was incubated with the resins on the rotating wheel for 1 hour at 4 °C. Finally, the flow through was discarded and the resins were washed 3 times with the dilution buffer, resuspended in SDS-loading buffer (50 mM Tris pH 6.8, 2% SDS, 5% β-mercaptoethanol, 10% glycerol, bromophenol blue) and boiled for 5 minutes at 95 °C.

### SDS-PAGE, Western blots, and Far-Western blots

Proteins were diluted in SDS-loading buffer and heated at 95 °C for 5 minutes. The samples were then run on Tris/glycine/SDS-polyacrylamide gel (the 12% and 18% separating gel for GST-tagged PHDs and histone proteins, respectively). For the Western blots, proteins were wet-transferred onto a nitrocellulose membrane (100 V, 1 hour, 4 °C). The membrane was blocked with 5% non-fat milk, incubated with a specific primary antibody for rabbit anti-histone H3 (Cell Signaling Technologies) or goat anti-GFP (R&D System) overnight at 4 °C. Next day, the membrane was washed 3 times with PBS-T (PBS with 0.1% Tween20), incubated at room temperature with anti-rabbit or anti-goat HRP-conjugated secondary antibodies (Sigma) for 1 hour and washed three times with PBS-T. For detection of GST-tagged proteins, after blocking with 5% non-fat milk, the membrane was incubated at room temperature with an anti-GST HRP-conjugated primary antibody (Abcam) for 1 hour and washed three times with PBS-T.

For Far-Western blotting, 5 μg of mononucleosomes were resolved using an 18% SDS-PAGE gel, wet-transferred onto a nitrocellulose membrane (1 hour, 100 V, 4 °C), and blocked for 1 hour at room temperature or overnight at 4 °C with 5% non-fat milk. The membranes were then washed three times with TBS-T and once with interaction buffer (20 mM HEPES 7.5, 100 mM KCl, 10% glycerol, 1 mM DTT, 150 μM ZnSO_4_). A subsequent incubation with 100 nM PHD domains in interaction buffer for 1 hour at room temperature was performed. The membranes were again washed three times with TBS-T and incubated with an anti-GST HRP-conjugated antibody (Abcam) for 1 hour at room temperature. Finally, three washes times with TBS-T was done.

Immunodetection was performed using self-made ECL (0.2 mM coumaric acid, 1.25 mM luminol, 100 mM Tris 8.5, 0.03% H_2_O_2_) or Femto ECL (ThermoScientific). The chemiluminescent signals were scanned from the membranes using a BioRad documentation system. All antibodies used are listed in the Table S3.

### Standard CUT&RUN (C&R) and greenCUT&RUN (gC&R)

For standard C&R [68], the protocol published by Janssens and Henikoff (dx.doi.org/10.17504/protocols.io.zcpf2vn) was used. For gC&R, the protocol described by Nizamuddin and colleagues [69], with minor modifications, was used. Briefly, in both protocols, 0.5 million HeLa S3 cells were collected by trypsinisation and washed twice with wash buffer (20 mM HEPES–KOH pH 7.5, 150 mM NaCl, 0.5 mM spermidine and EDTA-free complete protease inhibitor cocktail). In parallel, BioMag®Plus Concanavalin A (conA) magnetic beads (10 μl per 0.5 million HeLa S3 cells) were incubated twice with binding buffer (20 mM HEPES–KOH pH 7.5, 10 mM KCl, 1 mM CaCl_2_ and 1 mM MnCl_2_). The washed cells were then mixed with the conA beads and incubated on a vortexer for 8 minutes. All steps up to this point were performed at room temperature to minimise cellular stress and DNA fragmentation.

For C&R, conA immobilised cells were incubated with 100 μl antibody buffer (20 mM HEPES–KOH pH 7.5, 150 mM NaCl, 0.5 mM spermidine and EDTA-free protease inhibitor cocktail, 0.01% digitonin, 2 mM EDTA) supplemented with the antibody overnight on a rotating wheel at 4 °C. All antibodies used are listed in the Table S3.Following overnight incubation, the cells were washed twice with ice-cold digitonin buffer and incubated on a nutator for 1 hour with 50 μl digitonin buffer containing 1.5 μl C&R MNase (Cell Signalling Technologies). For gC&R, conA immobilised cells were incubated for 4 minutes at room temperature with antibody buffer and then washed twice with digitonin buffer. Next, the immobilised cells were incubated in 100 μl digitonin buffer with 0.4 ng of the fusion protein consisting of anti-GFP nanobody-MNase (a kind gift from Marc Timmers) for 1 hour at 4 °C on a rotating wheel.

In both protocols, cells were then washed two times with ice-cold digitonin buffer, resuspended in 100 μl digitonin buffer, and kept on a wet-ice bath for at least 5 minutes. Next, MNase digestion was activated by adding CaCl_2_ to a 2 mM final concentration. After 30 minutes, digestion was stopped by adding 100 μl 2x STOP buffer (340 mM NaCl, 20 mM EDTA, 10 mM EGTA, 0.01% digitonin, 100 μg/ml of RNase A [ThermoScientific]). Next, the mixtures were incubated in a thermoblock at 37 °C to release chromatin-MNase complexes. Supernatants (200 μl) were supplemented with 2 μl of 10% (w/v) SDS and 2.5 μl proteinase K (20 mg/ml; ThermoScientific), and incubated for 1 hour at 50 °C in a thermoblock. DNA was extracted using a ChIP DNA Clean & Concentrator kit (ZymoResearch). For background correction in the C&R experiments, rabbit normal IgG (Sigma-Aldrich) was used, and in the gC&R experiments, the parental HeLa S3 and NLS-EYFP negative control cell lines were used.

### Library preparation

For library preparation, a NEBNext II Ultra DNA Prep Kit was used according to Liu’s protocol (dx.doi.org/10.17504/protocols.io.bagaibse). Briefly, DNA isolated after C&R or gC&R was mixed with END Prep Enzyme Mix and Reaction Buffer, and incubated in a thermocycler at 20 °C for 30 minutes, and at 50 °C for 1 hour. Next, NEB Next Adaptor (25x diluted, 0.6 μM) was ligated using Ligation Master Mix and Ligation Enhancer for 15 minutes at 20 °C, followed by hairpin cleavage by USER enzyme for 15 minutes at 37 °C. For the cleaning of adaptor-ligated DNA, left-side selection using 1.7x AMPure XP beads (Beckman Coulter) was used. The purified DNA was mixed with Q5 MasterMix, Universal PCR primer, appropriate index primer (NEB Single Index, NEBNext® Multiplex Oligos for Illumina), and used for PCR in a thermocycler (BioRad, Eppendorf). Finally, the amplified DNA libraries were cleaned with AMPure XP beads (first 0.55x right-sided, then left-sided in total 1.15x). The quality of libraries was verified using TapeStation (Perlan, Agilent).

### NGS sequencing and data analysis

The concentration of the libraries was measured by qPCR using a Kapa Library Quantification kit (Kapa Biosciences, KK4824), according to the manufacturer’s protocol. Paired-end (2x 100 nt) sequencing (10 MR histone modifications, 30 MR full length hCOMPASS-like subunits and gC&R) was performed on an Illumina NovaSeq 6000 instrument, using a NovaSeq 6000 S1 Reagent Kit (200 cycles, Illumina) with the addition of 0.5% control library Phix (Illumina).

The upstream analysis was performed using the Galaxy server [70]. Original fastq files were trimmed using both the TrimGalore! and FastQTrimmer algorithms. The quality of the reads was confirmed by FastQC. Next, the paired-end reads were aligned to the human genome hg38 using Bowtie2 [71], filtered for minimum MAPQ20 and against a blacklist, and deduplicated. Peak calling was done using MACS2 [72,73]. Heatmaps were created using plotHeatmap from deeptools in Galaxy [74]. Reads were first analysed in IGV [75], then peaks genome-wide using KEGG [76] and GO [77] analysis on the Cistrome-GO webserver [78]. For the comparison of the bed files with HMM imputed data or histone marks, the standard Jaccard index [79], a measure of interval overlap from bedtools [80] was used. Alternatively, overlap was visualized using Venn diagram generated using Intervene [81].

### Conservation analysis (Shannon analysis)

For sequence conservation analysis, UniProt [82] was queried for proteins annotated as KMT2A, KMT2B, KMT2C, or KMT2D. The program cd-hit [83] was used to cluster sequences at 80% sequence similarity. Fusion proteins or truncated proteins (any proteins with a length less than 90% of the length of the reference human protein) were excluded from the analysis. To avoid bias from multiple annotated splice variants, only one protein per paralogue and species was retained. Proteins were aligned using Clustal Omega [84]. As in sequence logos, sequence conservation was defined as the difference between the maximally possible and actual Shannon entropy in bits (log2(20)+Σpilog2pi, where pi is the frequency of amino acid “i” in a given position). As scores were needed with regard to reference to the human protein and not to a gapped alignment, BALCONY was not used directly, and the calculation of Shannon scores was re-implemented in Python.

### Structural analysis based on AlphaFold structure predictions

For structural analysis of clustered PHD domains (2A13, 2C14, 2C57, 2D13 and 2D46), AlphaFold prediction results were used [40]. Based on the COSMIC data, Pymol and Chimera software was used for the labelling of the mutated sites.

### COSMIC data analysis

The data for the analysis of the distribution of mutations throughout the length of the KMT2A-D proteins and in the patient population was obtained from the Catalogue of Somatic Mutations in Cancer (COSMIC) release v.92 [39]. The COSMIC Mutation Data (Genome Screens) dataset containing data on mutations found in whole genome sequencing (WGS) analysis was downloaded from the database as a filtered file to comprise the changes found in the KMT2A–D genes in all the malignancies described in COSMIC. The data from targeted screens (non-WGS) was not used in the distribution analysis of mutations as too many data points (~92%) are missing in relation to the type and location of mutations for the non-WGS dataset to be informative.

For the analysis of the distribution of mutations among patients, the COSMIC Sample Features non-filtered dataset was downloaded to calculate the number of patients diagnosed with a given malignancy and to assign the mutations described in the Mutation Data file to individuals. The data points from the WGS Mutation Data dataset with ‘Mutation description’ assigned as ‘Unknown’ (33% of the whole dataset) were removed from the analysis. Finally, the data was filtered for the position within the reading domain.

## STATEMENTS AND DECLARATIONS

### Author contribution

Conceptualization of the study: ASC, MB. Manuscript preparation: ASC and MB with the input from the other authors. Preparation of figures: ASC, MB, MK, KM. Conservation and structural analysis: MB. Protein purification, Far-Western, GST pull-down, peptide pull-down, immunofluorescence, green/CUT&RUN, data deposition: ASC. Generation of HeLa S3 stable cell lines: MP. Cell sorting: KM. Optimisation of PHD expression constructs: AAK, MB, ASC. Bioinformatics analysis of CUT&RUN data: MB and ASC. Analysis of COSMIC data: MK, ASC, MB. Supervision of the study and funding: MB. All authors read and approved the final manuscript.

## Acknowledgement

We thank Marc Timmers (Freiburg University, Germany) for the kind gift of MNase-anti-GFP nanobody, Diagenode for antibodies against modified histones used in CUT&RUN experiments, Agnieszka Rawluszko-Wieczorek (Poznan University of Medical Sciences, Poland) for her help in cloning some plasmids, Dorota Adamska (CeNT, Warsaw University, Poland) for Illumina sequencing, Derek Janssens (Fred Hutchinson Cancer Research Center, USA) for advices regarding CUT&RUN protocols, Bartlomiej Czerwinski who helped with informatics and Karim Abu Nahia and Kamil Jastrzębski (IIMCB core facility) for their help. We are also grateful to Albert Jeltsch (Stuttgart University, Germany), Agnieszka Rawluszko-Wieczorek (Poznan University of Medical Sciences, Poland) and Aleksandra Pekowska (Nencki Institute of Experimental Biology, Warsaw, Poland) for the fruitful discussions, Anton Slyvka, Honorata Czapińska and Terry Karimi (International Institute of Molecular and Cell Biology) for comments on manuscript and to Katarzyna Szafran for comments on figures. NGS was performed at the Genomics Core Facility CeNT UW, using the NovaSeq 6000 platform financed by the Polish Ministry of Science and Higher Education (6817/IA/2018 2018-04-10).

## Data availability

All raw and processed sequencing green/CUT&RUN data generated in this study have been submitted to the NCBI Gene Expression Omnibus (GEO; https://www.ncbi.nlm.nih.gov/geo/) under accession number: GSE185921.

