## Supplementary information for "Clustered PHD domains in KMT2/MLL proteins are attracted by H3K4me3 and H3 acetylation-rich active promoters and enhancers"

### Supplementary methods

#### Immunofluorescence

Cells for immunofluorescence were seeded onto cover slides at the bottom of 24 well plates (50 000 cells per well). After 24 hours, the cells were washed twice with PBS, fixed for 10 minutes at room temperature (4% paraformaldehyde in PBS), and washed twice with PBS again. The cover slides were either used immediately or stored sealed with parafilm for up to two weeks at 4 °C. For immunostaining, cells were incubated with permeabilisation buffer (0.25% TritonX100 in PBS) for 20 minutes at room temperature, washed once with PBS, placed in blocking buffer (3% bovine serum albumin, BSA in PBS) for 30 minutes, incubated with primary antibody diluted in Ab buffer (1% BSA in PBS) in a humid box for 1 hour at room temperature or overnight at 4 °C, washed two times with PBS, incubated for 1 hour with secondary antibody and DAPI in Ab buffer, and washed two times with PBS. The cover slides were then briefly washed with MilliQ water and put on ethanol-cleaned glass slides containing Mowiol. The specimens were kept protected from light and analysed by confocal microscopy (LSM 800, Zeiss). All antibodies used are listed in the Table S3.

a

```
Human        1316  GG-AHGGRGRGRARLKSTASSIETLVVADIDSSPSKEEEEEDDDTMQNTVVLFSNTDKFV  1374
Cow          1281  GG-AHGGRGRGRARLKSTTSSIETLVVADIDSSPSKEEEDDDDDTMQNTVVLFSNTDKFV  1339
Pig           903  GG-AHGGRGRGRARLKSTTSSIETLVVADIDSSPSKEEEEDDDDTMQNTVVLFSNTDKFV   961
Quinea pig   1041  GG-AHGGRGRGRARLKSTTSSIETLVVADIDSSPSKEEEEEDDDTMQNTVVLFSNTDKFV  1099
Squirrel     1268  GG-AHGGRGRGRARLKSTTSSIETL-VADIDSSPSKEEEEEDDDTMQNTVVLFSNTDKFV  1325
Giant panda  1026  GG-AHGGRGRGRARLKSTTSSIETLVVADIDSSPSKEEEDDDDDTMQNTVVLFSNTDKFV  1084
Dog          1309  GG-AHGGRGRGRARLKSTTSSIETLVVADIDSSPSKEEEDDDDDTMQNTVVLFSNTDKFV  1367
Marmoset     1316  GG-AHGGRGRGRARLKSTASSIETLVVADIDSSPSKEEEEEDDDTMQNTVVLFSNTDKFV  1374
Mouse        1272  GGAAHGGRGRGRARLKSTTSSVETL-VADIDSSPSKEEEEEDDDTMQNTVVLFSNTDKFV  1330
                   ** ***************:**:*** *************::*******************

Human        1375  LMQDMCVVCGSFGRGAEGHLLACSQCSQCYHPYCVNSKITKVMLLKGWRCVECIVCEVCG  1434
Cow          1340  LMQDMCVVCGSFGRGAEGHLLACSQCSQCYHPYCVNSKITKVMLLKGWRCVECIVCEVCG  1399
Pig           962  LMQDMCVVCGSFGRGAEGHLLACSQCSQCYHPYCVNSKITKVMLLKGWRCVECIVCEVCG  1021
Quinea pig   1100  LMQDMCVVCGSFGRGAEGHLLACSQCSQCYHPYCVNSKITKVMLLKGWRCVECIVCEVCG  1159
Squirrel     1326  LMQDMCVVCGSFGRGAEGHLLACSQCSQCYHPYCVNSKITKVMLLKGWRCVECIVCEVCG  1385
Giant panda  1085  LMQDMCVVCGSFGRGAEGHLLACSQCSQCYHPYCVNSKITKVMLLKGWRCVECIVCEVCG  1144
Dog          1368  LMQDMCVVCGSFGRGAEGHLLACSQCSQCYHPYCVNSKITKVMLLKGWRCVECIVCEVCG  1427
Marmoset     1375  LMQDMCVVCGSFGRGAEGHLLACSQCSQCYHPYCVNSKITKVMLLKGWRCVECIVCEVCG  1434
Mouse        1331  LMQDMCVVCGSFGRGAEGHLLACSQCSQCYHPYCVNSKITKVMLLKGWRCVECIVCEVCG  1390
                   ************************************************************

Human        1435  QASDPSRLLLCDDCDISYHTYCLDPPLLTVPKGGWKCKWCVSCMQCGAASPGFHCEWQNS  1494
Cow          1400  QASDPSRLLLCDDCDISYHTYCLDPPLLTVPKGGWKCKWCVSCMQCGAASPGFHCEWQNS  1459
Pig          1022  QASDPSRLLLCDDCDISYHTYCLDPPLLTVPKGGWKCKWCVSCMQCGAASPGFHCEWQNS  1081
Quinea pig   1160  QASDPSRLLLCDDCDISYHTYCLDPPLLTVPKGGWKCKWCVSCMQCGAASPGFHCEWQNS  1219
Squirrel     1386  QASDPSRLLLCDDCDISYHTYCLDPPLLTVPKGGWKCKWCVSCMQCGAASPGFHCEWQNS  1445
Giant panda  1145  QASDPSRLLLCDDCDISYHTYCLDPPLLTVPKGGWKCKWCVSCMQCGAASPGFHCEWQNS  1204
Dog          1428  QASDPSRLLLCDDCDISYHTYCLDPPLLTVPKGGWKCKWCVSCMQCGAASPGFHCEWQNS  1487
Marmoset     1435  QASDPSRLLLCDDCDISYHTYCLDPPLLTVPKGGWKCKWCVSCMQCGAASPGFHCEWQNS  1494
Mouse        1391  QASDPSRLLLCDDCDISYHTYCLDPPLLTVPKGGWKCKWCVSCMQCGAASPGFHCEWQNS  1450
                   ************************************************************

Human        1495  YTHCGPCASLVTCPICHAPYVEEDLLIQCRHCERWMHAGCESLFTEDDVEQAADEGFDCV  1554
Cow          1460  YTHCGPCASLVTCPICHAPYVEEDLLIQCRHCERWMHAGCESLFTEDDVEQAADEGFDCV  1519
Pig          1082  YTHCGPCASLVTCPICHAPYVEEDLLIQCRHCERWMHAGCESLFTEDDVEQAADEGFDCV  1141
Quinea pig   1220  YTHCGPCASLVTCPICHAPYVEEDLLIQCRHCERWMHAGCESLFTEDDVEQAADEGFDCV  1279
Squirrel     1446  YTHCGPCASLVTCPICHAPYVEEDLLIQCRHCERWMHARCQSLLTEDDVEQAADEGFDCV  1505
Giant panda  1205  YTHCGPCASLVTCPICHAPYMEEDLLIQCRHCERWMHAGCESLFTEDDVEQAADEGFDCV  1264
Dog          1488  YTHCGPCASLVTCPICHTPYVEEDLLIQCRHCERWMHAGCESLFTEDDVEQAADEGFDCV  1547
Marmoset     1495  YTHCGPCASLVTCPICHAPYVEEDLLIQCRHCERWMHAGCESLFTEDDVEQAADEGFDCV  1554
Mouse        1451  YTHCGPCASLVTCPVCHAPYVEEDLLIQCRHCERWMHAGCESLFTEDEVEQAADEGFDCV  1510
                   **************:**:**:***************** *:**:***:************

Human        1555  SCQPYVVKPVAPVAPPELVPMKVKEPEPQYFRFEGVWLTETGMALLRNLTMSPLHKRRQR  1614
Cow          1520  SCQPYVVKPAAPVAPPELVPMKVKEPEPQYFRFEGVWLTETGMAVLRNLTMSPLHKRRQR  1579
Pig          1142  SCQPYVVKPAAPVAPPELVPMKVKEPEPQYFRFEGVWLTETGMAVLRNLTMSPLHKRRQR  1201
Quinea pig   1280  SCQPYVVKPVAPVAPPELVPMKVKEPEPQYFRFEGVWLTETGMAVLRNLTMSPLHKRRQR  1339
Squirrel     1506  SCQPYVVKPVVPVAPPELVPMKVKEPEPQYFRFEGVWLTETGMAVLRNLTMSPLHKRRQR  1565
Giant panda  1265  SCQPYVVKPAVPVAPPELVPMKVKEPEPQYFRFEGVWLTETGMAVLRNLTMSPLHKRRQR  1324
Dog          1548  SCQPYVVKPAAPVAPPELVPMKVKEPEPQYFRFEGVWLTETGMAVLRNLTMSPLHKRRQR  1607
Marmoset     1555  SCQPYVVKPVVPVAPPELVPMKVKEPEPQYFRFEGVWLTETGMAVLRNLTMSPLHKRRQR  1614
Mouse        1511  SCQPYVVKPVVPVAPPELVPVKVKEPEPQFFRFEGVWLTETGMAVLRNLTMSPLHKRRQR  1570
                   *********..*********:********:**************:***************
```

b

**IDENTITY**

| | KMT2A PHD1-3 | KMT2B PHD1-3 | KMT2C PHD1-4 | KMT2C PHD5-7 | KMT2D PHD1-3 |
|---|---|---|---|---|---|
| KMT2B PHD1-3 | 65% | | | | |
| KMT2C PHD1-4 | 32% | 32% | | | |
| KMT2C PHD5-7 | 31% | 34% | 36% | | |
| KMT2D PHD1-3 | 28% | 35% | 59% | 41% | |
| KMT2D PHD4-6 | 33% | 34% | 35% | 75% | 39% |

**SIMILARITY**

| | KMT2A PHD1-3 | KMT2B PHD1-3 | KMT2C PHD1-4 | KMT2C PHD5-7 | KMT2D PHD1-3 |
|---|---|---|---|---|---|
| KMT2B PHD1-3 | 80% | | | | |
| KMT2C PHD1-4 | 48% | 44% | | | |
| KMT2C PHD5-7 | 50% | 46% | 56% | | |
| KMT2D PHD1-3 | 37% | 41% | 73% | 56% | |
| KMT2D PHD4-6 | 49% | 46% | 53% | 84% | 55% |

**Supplementary Figure 1. Alignment of PHDs. a** MUSCLE alignment for 2D46 from selected mammals (human, cow, pig, guinea pig, squirrel, giant panda, dog, marmoset, and mouse). **b** Pairwise identity and similarity between clustered PHDs in human KMT2A-D proteins, based on blastp alignment.

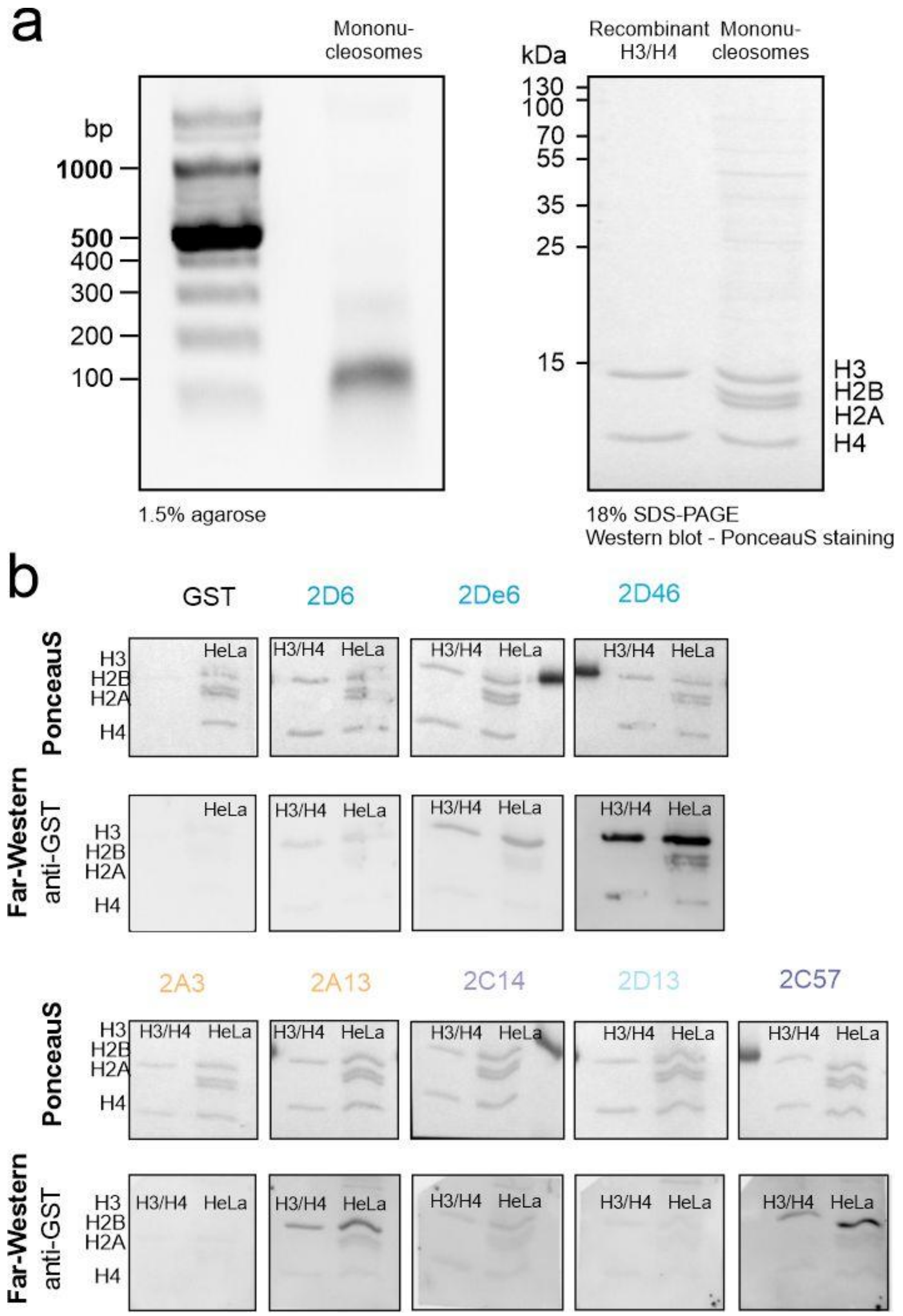


**Supplementary Figure 2. Biochemical characterisation of clustered PHDs binding histones.** **a** Characterization of isolated mononucleosomes from HeLa S3 cell line. **b** Far-Western of 100 nM singlets and clustered PHDs and recombinant H3/H4 or endogenous HeLa histones.

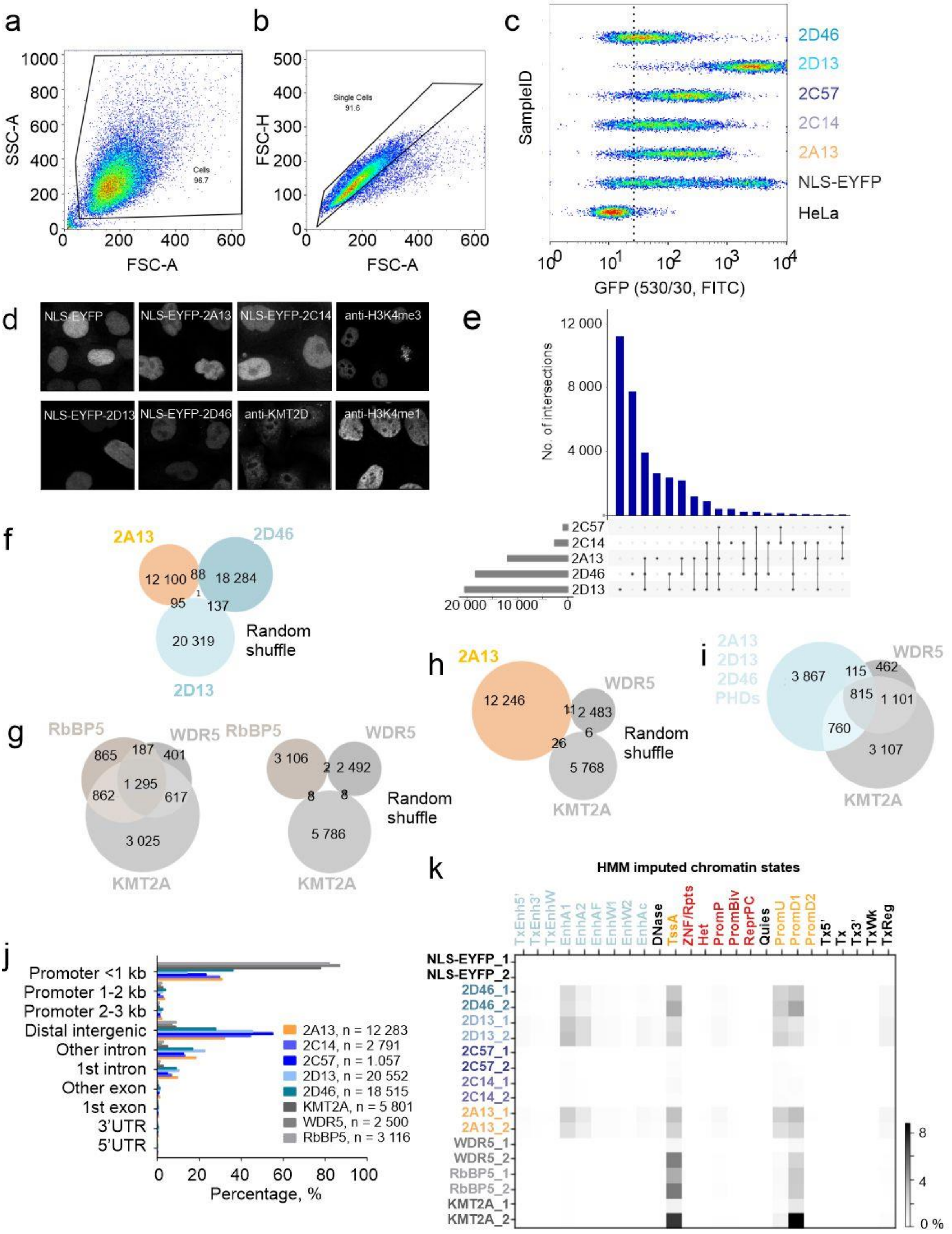

**Supplementary Figure 3. Characterisation of clustered PHDs in HeLa cells by greenCUT&RUN experiments. a-c** Flow cytometry analysis of Hela S3 cell lines. **a** Cells were gated based on Forward (FSC-Area) and Side Scatter (SSC-Area) parameters. **b** Cell doublets were discriminated from "Cells" based on Forward Scatter FSC-Height vs. Forward Scatter FSC-Area parameters. **c** EYFP fluorescence signal was detected using FITC (530/30) emission filter. For each cell line 5 000 cells were plotted. **d** Confocal microscopy of EYFP fluorescence in stable cell lines as well as immunofluorescence of KMT2D and of selected histone modifications (H3K4me1 and H3K4me3). **e** UpSet plot comparing peaks of all clustered PHDs. **f** Venn diagram comparing random shuffled peaks of the PHDs (2A13, 2D13, 2D46), **g** full-length hCOMPASS-like subunits (KMT2A, RbBP5, WDR5), **h** random shuffle of 2A13 PHDs and full-length WDR5 and KMT2A, **i** common peaks of PHDs (2A13, 2D13, 2D46) with full-length WDR5 and KMT2A. **j** Genomic distribution of binding sites of PHDs and hCOMPASS-like subunits as determined using MACS2. **k** Jaccard analysis comparing MACS2 bed files from single gC&R and C&R experiments with bed files for HMM (Hidden Markov Model) imputed chromatin states.

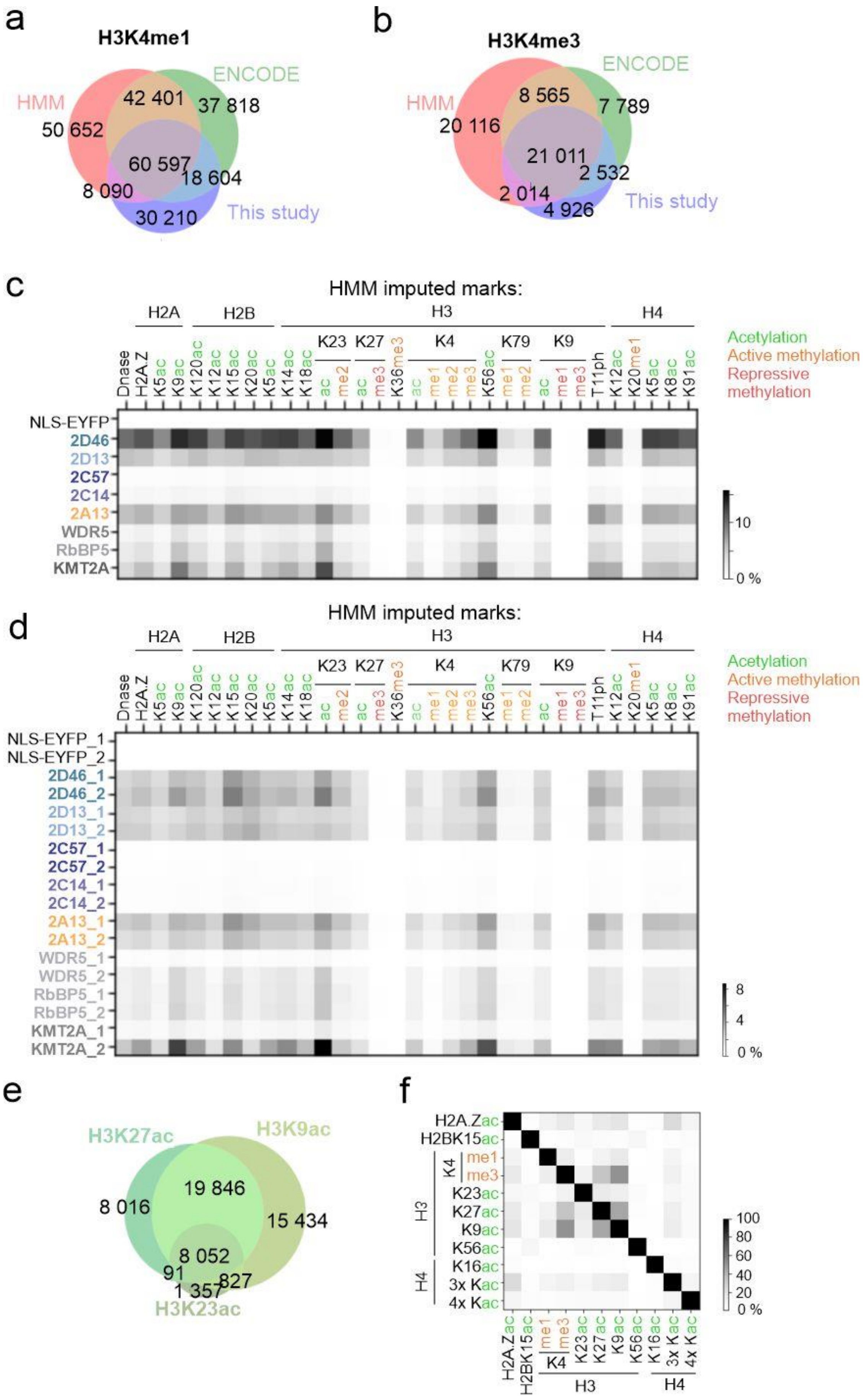

a
H3K4me1
HMM
ENCODE
This study
42 401
37 818
50 652
60 597
18 604
8 090
30 210
b
H3K4me3
HMM
ENCODE
This study
8 565
7 789
20 116
21 011
2 532
2 014
4 926
c
HMM imputed marks:
H2A
H2B
H3
H4
K23
K27
K4
K79
K9
Dnase
H2A.Z
K5ac
K9ac
K120ac
K12ac
K15ac
K20ac
K5ac
K14ac
K18ac
ac
me2
ac
me3
K36me3
ac
me1
me2
me3
K56ac
me1
me2
ac
me1
me3
T11ph
K12ac
K20me1
K5ac
K8ac
K91ac
Acetylation
Active methylation
Repressive methylation
NLS-EYFP
2D46
2D13
2C57
2C14
2A13
WDR5
RbBP5
KMT2A
10
0 %
d
HMM imputed marks:
NLS-EYFP_1
NLS-EYFP_2
2D46_1
2D46_2
2D13_1
2D13_2
2C57_1
2C57_2
2C14_1
2C14_2
2A13_1
2A13_2
WDR5_1
WDR5_2
RbBP5_1
RbBP5_2
KMT2A_1
KMT2A_2
8
4
0 %
e
H3K27ac
H3K9ac
H3K23ac
8 016
19 846
15 434
8 052
91
1 357
827
f
H2A.Zac
H2BK15ac
K4
me1
me3
K23ac
K27ac
K9ac
K56ac
K16ac
3x Kac
4x Kac
H3
H4
100
80
60
40
20
0 %

**Supplementary Figure 4. Characterization of the investigated chromatin marks in HeLa S3 cell lines. a-b** Venn diagrams comparing **a** H3K4me1 or **b** H3K4me3 from our CUT&RUN experiments with ENCODE ChIP-seq and HMM imputed data. **c** Jaccard analysis comparing MACS2 peaks of PHDs and full-length hCOMPASS-like subunits (KMT2A, Wdr5, RbBP5) with HMM imputed chromatin marks with merged replicas and **d** with single replicas. **e** Venn diagrams comparing the overlap of chromatin marks most associated with PHD domain binding (H3K4me1/me3, H3K9ac, H3K27ac, H3K23ac and H2BK15ac). **f** Jaccard analysis showing the overlap between chromatin marks analysed by CUT&RUN.

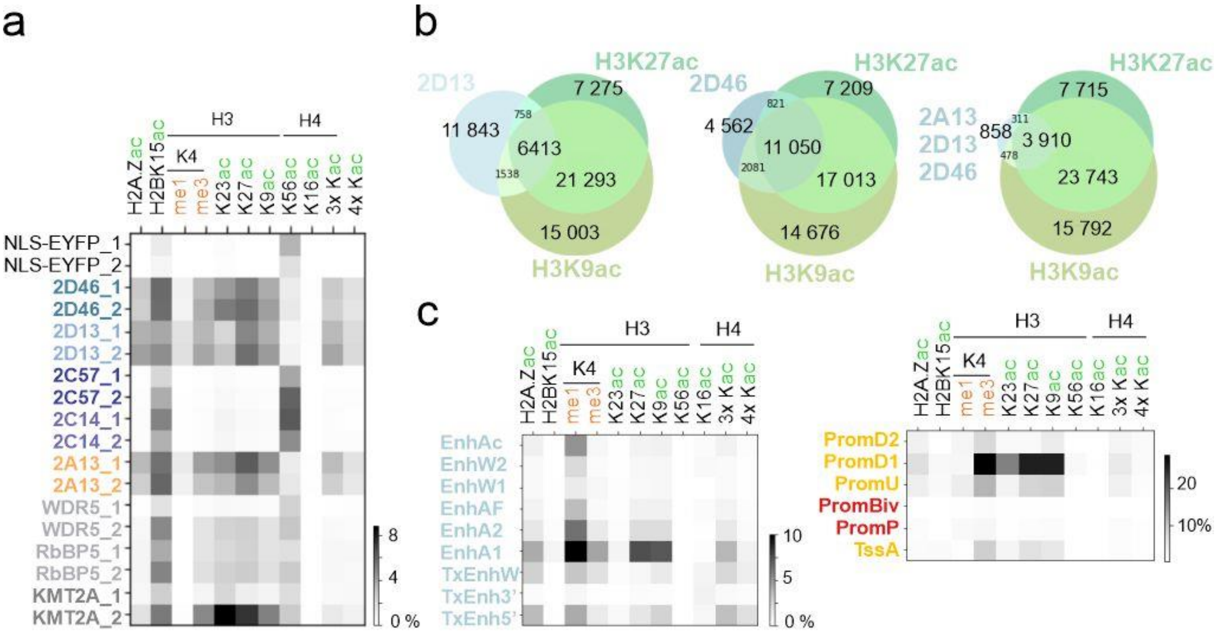


**Supplementary Figure 5. Chromatin marks at binding sites of PHDs. a** Jaccard analysis of PHDs binding sites and histone modifications, without replica merging. **b** Venn diagrams comparing the MACS2 peaks of selected PHDs (2D13, 2D46) with the strongest histone marks (H3K9ac, H3K27ac). **c** Histone marks at enhancer (left) and promoter (right) imputed chromatin states.

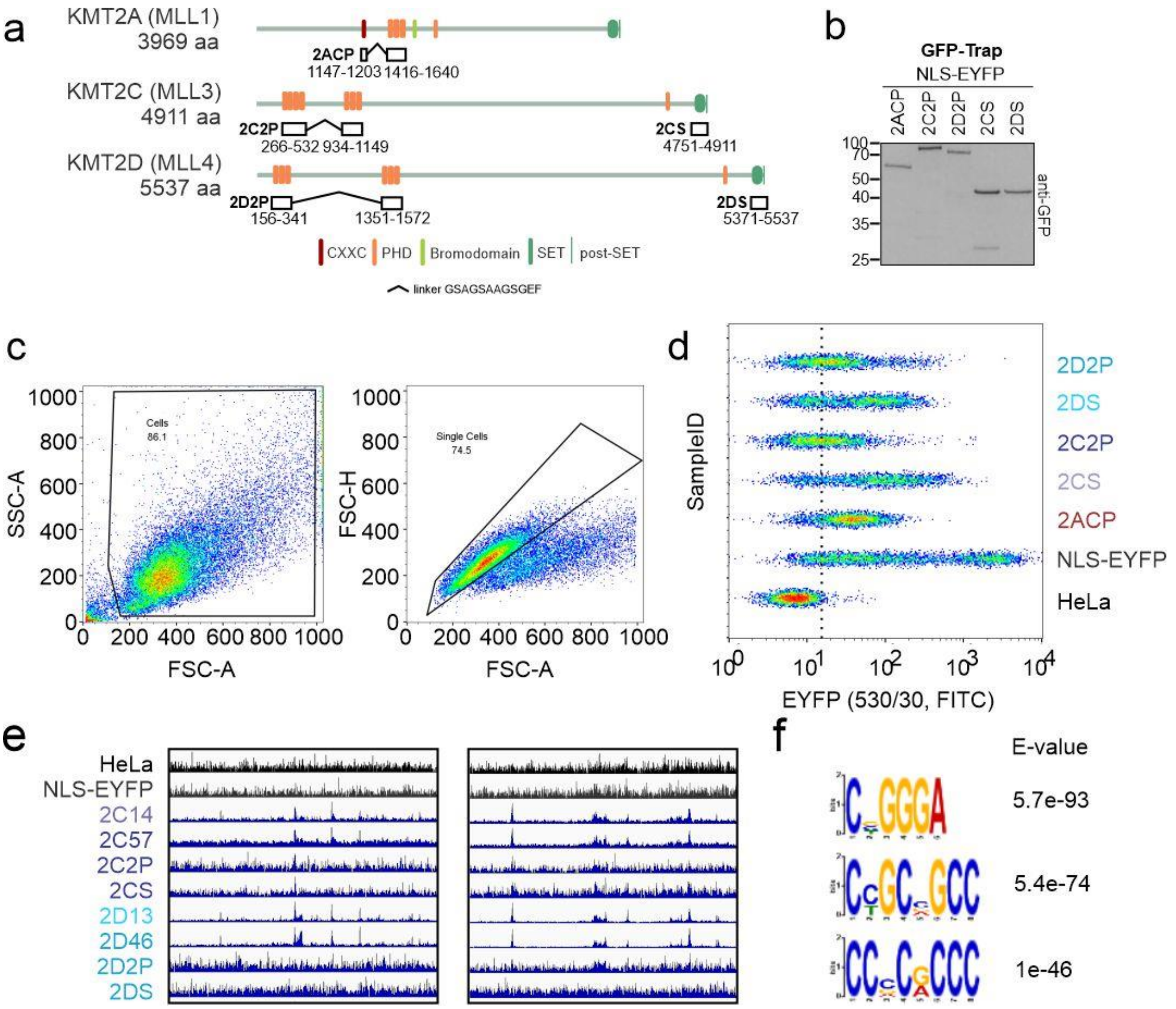


**Supplementary Fig. 6** **a** Tandem reader domain constructs. 2ACP – fusion of CXXC and PHDs from KMT2A. 2C2P/2D2P – tandem of two clustered PHDs from KMT2C or KMT2D proteins, respectively. 2CS/2DS – SET domain of KMT2C or KMT2D proteins, respectively. **b** Western blot analysis with anti-GFP antibody after GFP-Trap enrichment of stable HeLa cell line lysates. **c-d** Flow cytometry analysis of Hela S3 stable cell lines. **c** Left panel - cells were gated based on Forward (FSC-Area) and Side Scatter (SSC-Area) parameters. Right panel **-** Cell doublets were discriminated from "Cells" based on Forward Scatter FSC-Height vs. Forward Scatter FSC-Area parameters. **d** EYFP fluorescence signal was detected using FITC (530/30) emission filter. For each cell line 5 000 cells were plotted. **e** IGV browser snapshot showing the reads of greenCUT&RUN for tandem 2C2P/2D2P and 2CS/2DS stable cell lines. **f** Top 3 hits of the MEME-ChIP DNA motif analysis for 2ACP binding sites as determined by MACS2.

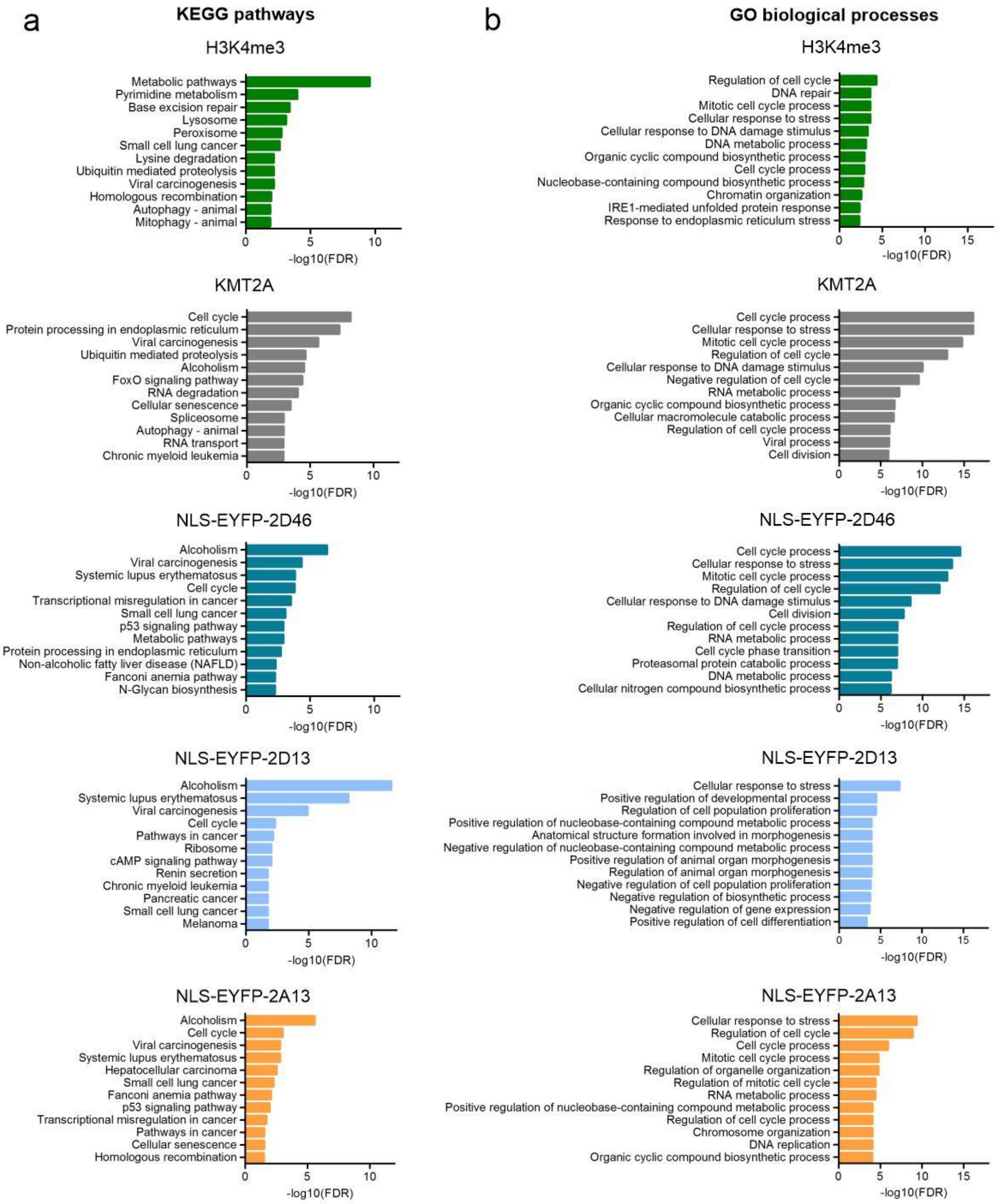


**Supplementary Figure 7. Characterisation of the clustered PHDs and full-length KMT2A binding sites.** **a** Top twelve KEGG pathways (with the lowest FDR) of KMT2A and selected clustered PHDs (2A13 2D13, 2D46). **b** Top twelve GO biological processes (with the lowest FDR) of KMT2A and selected clustered PHDs (2A13, 2D13, 2D46).

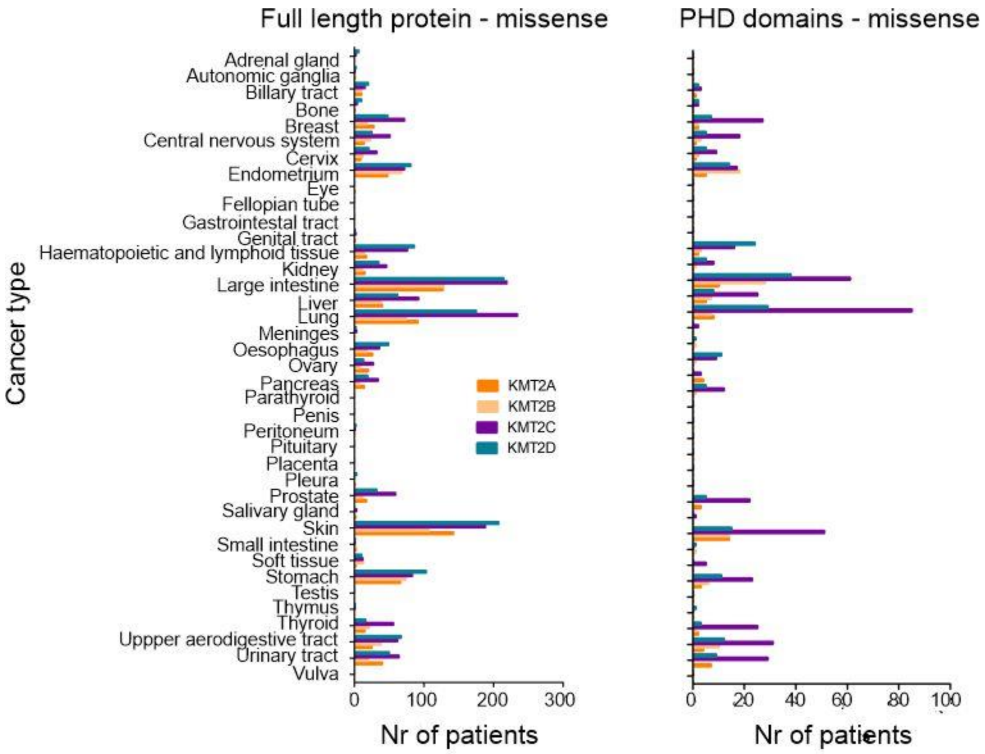


**Supplementary Figure 8. Distribution of missense mutations in the cancer patients, analyzed for full-length proteins or PHD domains only.**

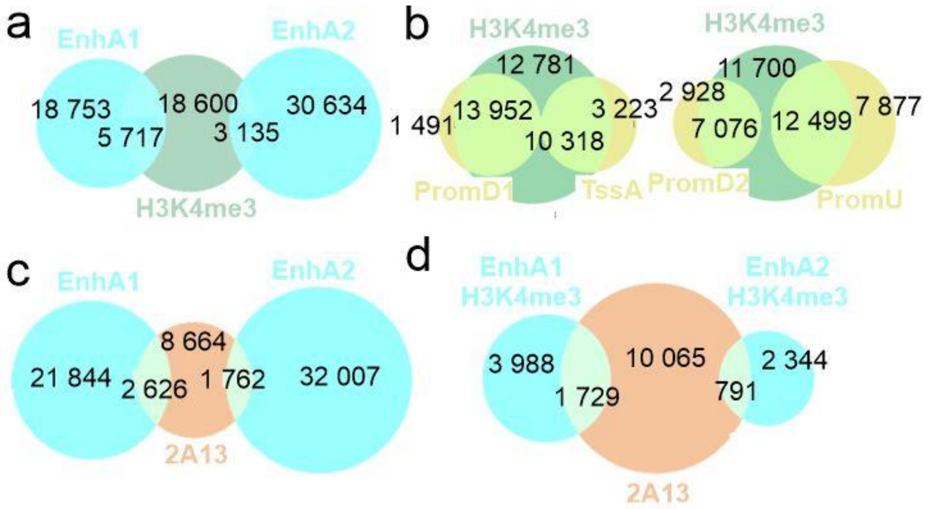


**Supplementary Figure 9. H3K4me3 and 2A13 at active promoters and enhancers.** Venn diagrams comparing peaks of H3K4me3 with imputed active **a** enhancers EnhA1, EnhA2, **b** promoters TssA and PromD1 (left panel), PromD2 and PromU (right panel), **c** 2A13 PHDs with imputed enhancers EnhA1/2 and **d** fraction of these enhancers decorated with H3K4me3.

**Tabel S1. Synthetic genes used in this study.**

| OLIGONUCLEOTIDE SEQUENCE | SOURCE | IDENTIFIER |
|---|---|---|
| **Synthetic genes for recombinant proteins** | | |
| TATAAAAATAGGCGTATCACGAGGCATATGGGATCCAAA<br>GGTAATTTTTGTCCGCTGTGCGATAAATGCTATGATGATG<br>ACGATTACGAGAGCAAAATGATGCAGTGTGGTAAATGTG<br>ATCGTTGGGTTCATAGCAAATGCGAAAATCTGTCCGATG<br>AGATGTATGAAATTCTGAGCAATCTGCCGGAAAGCGTTG<br>CATATACCTGTGTTAATTGTACCGAACGTCATCCGGCAG<br>AATAACTCGAGGCGGCCGCCCGCTGAGCAATAACTAGCA | ThermoScientific | 2A3 |
| TATAAAAATAGGCGTATCACGAGGCATATGGGATCCGAA<br>ATGGGTGGTCTGGGTATTCTGACCAGCGTTCCGATTACAC<br>CGCGTGTTGTTTGTTTTCTGTGTGCAAGCAGCGGTCATGT<br>TGAATTTGTTTATTGTCAGGTTTGCTGCGAACCGTTTCAC<br>AAATTTTGTCTGGAAGAAAATGAACGTCCGCTGGAAGAT<br>CAGCTGGAAAATTGGTGTTGTCGTCGTTGTAAATTTTGCC<br>ATGTTTGTGGTCGTCAGCATCAGGCAACCAAACAGCTGC<br>TGGAATGTAATAAATGCCGCAATAGCTATCATCCGGAAT<br>GTCTGGGTCCGAATTATCCGACCAAACCGACAAAAAAGA<br>AAAAGGTTTGGATCTGCACCAAATGCGTTCGTTGCAAAA<br>GCTGTGGTAGCACCACACCTGGTAAAGGTTGGGATGCAC<br>AGTGGTCACATGATTTTAGCCTGTGTCATGATTGTGCAAA<br>ACTGTTTGCCAAAGGCAATTTTTGTCCGCTGTGTGATAAA<br>TGCTATGATGATGACGATTACGAAAGCAAGATGATGCAG<br>TGTGGTAAATGTGATCGTTGGGTTCATAGCAAATGCGAA<br>AATCTGTCCGATGAGATGTATGAAATTCTGAGCAATCTG<br>CCGGAAAGCGTTGCATATACCTGTGTTAATTGTACCGAA<br>CGTCATCCGGCAGAATGGCGTCTGGCACTGGAAAAAGAA<br>CTGTAACTCGAGGCGGCCGCCCGCTGAGCAATAACTAGC<br>A | ThermoScientific | 2A13 |
| TATAAAAATAGGCGTATCACGAGGCATATGGGATCCGAA<br>GAACCGCTGCTGGTTAATGTTGATAAAGCAGTTGTTAGC<br>GGTAGCACCGAACGTTGTGCATTTTGTAAACATCTGGGT<br>GCCACCATTAAATGCTGTGAAGAAAAATGTACGCAGATG<br>TATCATTATCCGTGTGCAGCCGGTGCAGGCACCTTTCAGG<br>ATTTTAGCCATATTTTTCTGCTGTGTCCGGAACATATTGA<br>TCAGGCACCGGAACGTAGCAAAGAAGATGCAAATTGCG<br>CAGTTTGTGATAGTCCGGGTGATCTGCTGGATCAGTTTTT<br>TTGTACCACCTGTGGTCAGCATTATCATGGTATGTGTCTG<br>GATATTGCAGTTACACCGCTGAAACGTGCAGGTTGGCAG<br>TGTCCTGAATGTAAAGTTTGTCAGAATTGTAAACAGAGC<br>GGTGAGGATAGCAAAATGCTGGTTTGCGATACCTGTGAT<br>AAAGGCTATCACACCTTTTGTCTGCAGCCTGTTATGAAAA<br>GCGTTCCGACCAATGGTTGGAAATGTAAAAATTGCCGCA<br>TTTGCATTGAATGCGGCACCCGTAGCAGCAGCCAGTGGC<br>ATCATAATTGTCTGATTTGCGATAATTGCTATCAGCAGCA<br>GGATAATCTGTGTCCGTTTTGTGGTAAATGTTATCATCCG<br>GAACTGCAGAAAGACATGCTGCATTGTAATATGTGTAAA<br>CGTTGGGTTCATCTGGAATGTGATAAACCGACCGATCAT<br>GAACTGGATACCCAGCTGAAAGAAGAATATATCTGCATG<br>TATTGCAAACACCTGGGAGCAGAAATGGATCGTCTGCAA | ThermoScientific | 2C14 |

| | | |
|---|---|---|
| CCGGGTGAAGAATAACTCGAGGCGGCCGCCCGCTGAGCA ATAACTAGCA | | |
| TATAAAAATAGGCGTATCACGAGGCATATGGGATCCAGC CTGAGCAGCTGTCCGGTTTGTTATCGTAATTATCGTGAAG AGGATCTGATTCTGCAGTGTCGTCAGTGTGATCGTTGGAT GCATGCAGTTTGTCAGAATCTGAATACCGAAGAAGAAGT TGAAAACGTTGCCGATATTGGTTTTGATTGTAGCATGTGT CGTCCGTATATGCCTGCAAGCTAACTCGAGGCGGCCGCC CGCTGAGCAATAACTAGCA | ThermoScientific | 2C7 |
| TATAAAAATAGGCGTATCACGAGGCATATGGGATCCGTT TGGTGTCGTCATTGTGGTGCAACCAGCGCAGGTCTGCGTT GTGAATGGCAGAATAACTATACCCAGTGTGCACCGTGTG CAAGCCTGAGCAGCTGTCCGGTTTGTTATCGTAATTATCG TGAAGAGGATCTGATTCTGCAGTGTCGTCAGTGCGATCG TTGGATGCATGCAGTTTGTCAGAATCTGAATACCGAAGA AGAAGTTGAAAACGTTGCCGATATTGGTTTTGATTGTAG CATGTGTCGTCCGTATATGCCTGCAAGCAATTAACTCGA GGCGGCCGCCCGCTGAGCAATAACTAGCA | ThermoScientific | 2Ce7 |
| TATAAAAATAGGCGTATCACGAGGCATATGGGATCCACC GTTTGTGAAGCATGTGGTAAAGCAACCGATCCGGGTCGT CTGCTGCTGTGTGATGATTGTGATATTAGCTATCACACCT ATTGTCTGGACCCTCCGCTGCAGACCGTTCCGAAAGGTG GTTGGAAATGTAAATGGTGTGTTTGGTGTCGTCATTGTGG TGCAACCAGCGCAGGTCTGCGTTGTGAATGGCAGAATAA CTATACCCAGTGTGCACCGTGTGCAAGCCTGAGCAGCTG TCCGGTTTGTTATCGTAATTATCGTGAAGAGGATCTGATT CTGCAGTGTCGTCAGTGCGATCGTTGGATGCATGCAGTTT GTCAGAATCTGAATACCGAAGAAGAAGTTGAAAACGTTG CCGATATTGGTTTTGATTGTAGCATGTGTCGTCCGTATAT GCCTGCAAGCAATTAACTCGAGGCGGCCGCCCGCTGAGC AATAACTAGCA | ThermoScientific | 2C67 |
| TATAAAAATAGGCGTATCACGAGGCATATGGGATCCGAT GATGAAGAAAATAGCATGCATAATACCGTGGTTCTGTTT AGCAGCAGCGATAAATTCACCCTGAATCAGGATATGTGT GTTGTTTGTGGTAGCTTTGGTCAGGGTGCCGAAGGTCGTC TGCTGGCATGTAGCCAGTGTGGTCAGTGTTATCATCCGTA TTGTGTGAGCATCAAAATCACCAAAGTTGTGCTGAGCAA AGGTTGGCGTTGTCTGGAATGTACCGTTTGTGAAGCATGT GGTAAAGCAACCGATCCGGGTCGCCTGCTGCTGTGTGAT GATTGTGATATTAGCTATCACACCTATTGTCTGGACCCTC CGCTGCAGACCGTTCCGAAAGGTGGTTGGAAATGTAAAT GGTGTGTTTGGTGTCGTCATTGTGGTGCAACCAGCGCAG GTCTGCGTTGTGAATGGCAGAATAACTATACCCAGTGTG CACCGTGTGCAAGCCTGAGCAGCTGTCCGGTTTGTTATCG TAATTATCGTGAAGAGGATCTGATTCTGCAGTGCCGTCA GTGTGATCGTTGGATGCATGCAGTTTGTCAGAATCTGAAT ACCGAAGAAGAAGTTGAAAACGTTGCCGATATTGGTTTT GATTGTAGCATGTGTCGTCCGTATATGCCTGCAAGCAAT GTTCCGAGCAGCGATTAACTCGAGGCGGCCGCCCGCTGA GCAATAACTAGCA | ThermoScientific | 2C57 |
| TATAAAAATAGGCGTATCACGAGGCATATGGGATCCGAA CTGTGTGGTGTTGATAAAGCAATTTTTAGCGGTATTAGCC | ThermoScientific | 2D13 |

| | | |
|---|---|---|
| AGCGTTGTAGCCATTGTACCCGTCTGGGTGCAAGCATTCC GTGTCGTAGTCCGGGTTGTCCGCGTCTGTATCATTTTCCG TGTGCAACCGCAAGCGGTAGCTTTCTGAGCATGAAAACC CTGCAGCTGCTGTGTCCGGAACATAGCGAAGGTGCAGCA TATCTGGAAGAAGCACGTTGCGCAGTTTGTGAAGGTCCG GGTGAACTGTGCGACCTGTTTTTTTGTACCAGCTGTGGTC ATCATTATCATGGTGCATGTCTGGATACCGCACTGACCGC ACGTAAACGTGCAGGTTGGCAGTGTCCTGAATGTAAAGT TTGTCAGGCATGTCGTAAACCGGGTAATGATAGCAAAAT GCTGGTTTGTGAAACCTGCGATAAAGGCTATCATACCTTT TGTCTGAAACCGCCTATGGAAGAACTGCCTGCACATAGC TGGAAATGTAAAGCATGTCGTGTTTGTCGTGCATGTGGT GCAGGTAGCGCAGAACTGAATCCGAATAGCGAATGGTTT GAATAACTCGAGGCGGCCGCCCGCTGAGCAATAACTAGC A | | |
| TATAAAAATAGGCGTATCACGAGGCATATGGGATCCAGC CTGGTTACCTGTCCGATTTGTCATGCACCGTATGTTGAAG AGGATCTGCTGATTCAGTGTCGTCATTGTGAACGTTGGAT GCATGCAGGTTGTGAAAGCCTGTTTACCGAAGATGATGT TGAACAGGCAGCAGATGAAGGTTTTGATTGTGTTAGCTG TCAGCCGTATGTGGTGAAATAACTCGAGGCGGCCGCCCG CTGAGCAATAACTAGCA | ThermoScientific | 2D6 |
| TATAAAAATAGGCGTATCACGAGGCATATGGGATCCGTT AGCTGTATGCAGTGTGGTGCAGCAAGTCCGGGTTTTCATT GTGAATGGCAGAATAGCTATACCCATTGTGGTCCGTGTG CAAGCCTGGTTACCTGTCCGATTTGTCATGCACCGTATGT TGAAGAGGATCTGCTGATTCAGTGTCGTCATTGCGAACG TTGGATGCATGCAGGTTGTGAAAGTCTGTTTACCGAAGA TGATGTTGAACAGGCAGCAGATGAAGGTTTTGATTGTGT TAGCTGTCAGCCGTATGTGGTTAAACCGGTGTAACTCGA GGCGGCCGCCCGCTGAGCAATAACTAGCA | ThermoScientific | 2De6 |
| TATAAAAATAGGCGTATCACGAGGCATATGGGATCCGAA GAAGAGGAAGAGGACGACGATACCATGCAGAATACCGT TGTTCTGTTTAGCAACACCGATAAATTTGTGCTGATGCAG GATATGTGTGTTGTTTGTGGTAGCTTTGGTCGTGGTGCCG AAGGTCATCTGCTGGCATGTAGCCAGTGCAGCCAGTGTT ATCATCCGTATTGTGTGAATAGCAAAATCACCAAAGTGA TGCTGCTGAAAGGTTGGCGTTGTGTTGAATGTATTGTTTG TGAAGTTTGTGGTCAGGCAAGCGATCCGAGCCGTCTGCT GCTGTGTGATGATTGTGATATTAGCTATCACACCTATTGT CTGGACCCTCCGCTGCTGACCGTTCCGAAAGGTGGTTGG AAATGTAAATGGTGTGTTAGCTGTATGCAGTGTGGTGCA GCAAGTCCGGGTTTTCATTGTGAATGGCAGAATAGCTAT ACCCATTGTGGTCCGTGTGCAAGCCTGGTTACCTGTCCGA TTTGTCATGCACCGTATGTTGAAGAGGATCTGCTGATTCA GTGTCGTCATTGCGAACGTTGGATGCATGCAGGTTGTGA AAGTCTGTTTACCGAAGATGATGTTGAACAGGCAGCCGA TGAAGGTTTTGATTGCGTTAGCTGTCAGCCGTATGTGGTT AAACCGGTTGCACCGGTGGCACCGCCTGAACTGTAACTC GAGGCGGCCGCCCGCTGAGCAATAACTAGCA | ThermoScientific | 2D46 |
| **Synthethic genes for cellular expression** | | |
| TATAAAAATAGGCGTCTTAAGGCTAGCCGCCACCATGGC | ThermoScientific | 2A13_cellular |

| | | |
|---|---|---|
| TCCTAAGAAGAAGCGTAAGGTAGGTACCTCACCGGTAGG<br>CATATGTGTACAAGAGGAATGTCGACGAGATGGGCGGCC<br>TGGGCATCCTGACATCTGTGCCCATCACTCCCAGAGTCGT<br>GTGCTTCCTGTGTGCCTCTTCTGGCCACGTGGAATTCGTG<br>TACTGCCAAGTGTGCTGCGAGCCCTTCCACAAGTTCTGCC<br>TGGAAGAGAACGAGCGGCCCCTGGAAGATCAGCTGGAA<br>AACTGGTGCTGCCGGCGGTGCAAGTTCTGTCATGTCTGTG<br>GCAGACAGCACCAGGCCACCAAACAGCTGCTGGAATGC<br>AACAAGTGCCGGAACAGCTATCACCCCGAGTGTCTGGGC<br>CCTAACTACCCCACCAAGCCTACCAAGAAAAAGAAAGTG<br>TGGATCTGCACCAAATGCGTCCGGTGCAAGAGCTGCGGC<br>TCTACCACACCTGGCAAAGGCTGGGATGCTCAGTGGTCC<br>CACGATTTCAGCCTGTGCCACGATTGCGCCAAGCTGTTCG<br>CCAAGGGCAACTTCTGCCCTCTGTGCGACAAGTGCTACG<br>ACGACGATGACTACGAGAGCAAGATGATGCAGTGCGGC<br>AAGTGCGACAGATGGGTGCACTCCAAGTGCGAGAATCTG<br>AGCGGCACCGAGGACGAGATGTACGAGATCCTGAGCAA<br>CCTGCCTGAGAGCGTGGCCTACACCTGTGTGAACTGCAC<br>CGAAAGACACCCTGCCGAATGGCGGCTGGCCCTGGAAAA<br>AGAACTTTAAGGATCCCTCGAGGCGGCCGCCCGCTGAGC<br>AATAACTAGCA | | |
| TACCTCACCGGTAGGCATATGTGTACAAGAGGAATGTCG<br>ACGAGCCCCTGCTGGTCAACGTGGACAAAGCTGTGGTGT<br>CTGGCTCTACCGAGAGATGCGCCTTCTGCAAACACCTGG<br>GCGCCACCATCAAGTGCTGCGAAGAGAAGTGCACCCAGA<br>TGTATCACTACCCCTGTGCCGCTGGCGCCGGAACCTTTCA<br>GGATTTCAGCCACATCTTTCTGCTGTGCCCCGAGCACATC<br>GATCAGGCCCCTGAGAGAAGCAAAGAGGACGCCAATTG<br>CGCCGTGTGCGATAGCCCTGGCGATCTGCTGGATCAGTT<br>CTTCTGCACCACCTGTGGCCAGCACTACCACGGCATGTG<br>CCTGGATATTGCCGTGACACCCCTGAAGCGAGCCGGATG<br>GCAATGCCCTGAGTGCAAAGTGTGCCAGAACTGCAAGCA<br>GAGCGGCGAGGACAGCAAGATGCTCGTGTGCGACACCTG<br>TGACAAGGGCTACCACACCTTCTGCCTGCAACCTGTGAT<br>GAAGTCCGTGCCTACCAACGGCTGGAAGTGCAAGAACTG<br>CCGGATCTGCATCGAGTGCGGCACCAGATCTAGCAGCCA<br>GTGGCACCACAATTGCCTGATCTGCGACAACTGCTACCA<br>GCAGCAGGACAATCTGTGCCCCTTCTGCGGCAAGTGCTA<br>TCACCCCGAGCTGCAGAAAGACATGCTGCACTGCAACAT<br>GTGCAAGAGATGGGTGCACCTGGAATGCGACAAGCCCAC<br>CGATCACGAGCTGGACACCCAGCTGAAAGAAGAGTACAT<br>CTGCATGTATTGCAAGCACCTCGGAGCCGAGATGGACAG<br>ACTGCAGCCTGGCGAAGAATGAGGATCCCTCGAGGCGGC<br>CGCCCGCTGAGCAATAACTAGCA | ThermoScientific | 2C14_cellular |
| TACCTCACCGGTAGGCATATGTGTACAAGAGGAATGTCG<br>ACGACGACGAAGAGAACAGCATGCACAACACCGTGGTG<br>CTGTTCAGCAGCAGCGACAAGTTCACCCTGAACCAGGAT<br>ATGTGCGTCGTGTGCGGCTCTTTTGGACAGGGCGCTGAA<br>GGCAGACTGCTGGCCTGTTCTCAGTGCGGCCAGTGCTAT<br>CACCCTTACTGCGTGTCCATCAAGATCACCAAGGTGGTG<br>CTGAGCAAAGGCTGGCGGTGCCTGGAATGCACAGTGTGT<br>GAAGCCTGCGGCAAGGCCACAGATCCTGGTAGACTGCTG | ThermoScientific | 2C57_cellular |

| | | |
|---|---|---|
| CTGTGCGACGACTGCGACATCAGCTACCACACCTACTGT<br>CTGGACCCTCCACTGCAGACCGTGCCTAAAGGCGGATGG<br>AAGTGCAAGTGGTGCGTGTGGTGTAGACACTGCGGAGCC<br>ACATCTGCCGGACTGAGATGCGAGTGGCAGAACAACTAC<br>ACCCAGTGCGCCCCTTGTGCCAGCCTGAGTTCTTGCCCTG<br>TGTGCTACCGGAACTACAGAGAAGAGGACCTGATCCTGC<br>AGTGCCGGCAGTGCGATAGATGGATGCATGCCGTGTGCC<br>AGAACCTGAACACCGAGGAAGAGGTGGAAAACGTCGCC<br>GACATCGGCTTCGACTGCAGCATGTGCAGACCCTACATG<br>CCCGCCAGCAATGTGCCTAGCTCTGACTAAGGATCCCTC<br>GAGGCGGCCGCCCGCTGAGCAATAACTAGCA | | |
| TACCTCACCGGTAGGCATATGTGTACAAGAGGAATGTCG<br>ACGAGCTGTGCGGCGTGGACAAGGCCATCTTTAGCGGCA<br>TCAGCCAGAGATGCAGCCACTGCACAAGACTGGGCGCCA<br>GCATTCCTTGTAGAAGCCCTGGCTGCCCCAGACTGTATCA<br>CTTCCCTTGTGCTACAGCCAGCGGCAGCTTCCTGAGCATG<br>AAGACACTGCAGCTGCTGTGCCCTGAGCACTCTGAAGGC<br>GCCGCTTATCTGGAAGAGGCCAGATGCGCTGTGTGTGAA<br>GGACCTGGCGAGCTGTGTGACCTGTTCTTCTGTACCAGCT<br>GCGGCCACCACTATCACGGCGCCTGTCTGGATACAGCCC<br>TGACCGCCAGAAAACGCGCTGGATGGCAATGCCCCGAGT<br>GCAAAGTTTGCCAGGCCTGCAGAAAGCCCGGCAACGACT<br>CTAAGATGCTCGTGTGCGAGACATGCGACAAGGGCTACC<br>ACACCTTCTGCCTGAAGCCTCCAATGGAAGAACTGCCCG<br>CTCACAGCTGGAAGTGCAAGGCCTGTAGAGTGTGCAGAG<br>CCTGTGGCGCCGGATCTGCCGAGCTGAATCCTAACAGCG<br>AGTGGTTCGAATAAGGATCCCTCGAGGCGGCCGCCCGCT<br>GAGCAATAACTAGCA | ThermoScientific | 2D13_cellular |
| TACCTCACCGGTAGGCATATGTGTACAAGAGGAATGTCG<br>ACGAGGAAGAAGAAGAGGACGACGATACCATGCAGAAC<br>ACCGTGGTGCTGTTCAGCAACACCGACAAGTTCGTGCTG<br>ATGCAGGATATGTGCGTCGTGTGCGGCAGCTTTGGAAGA<br>GGCGCTGAAGGACATCTGCTGGCCTGTAGCCAGTGCTCC<br>CAGTGCTATCACCCCTACTGCGTGAACAGCAAGATCACC<br>AAAGTGATGCTGCTGAAAGGCTGGCGCTGCGTGGAATGC<br>ATCGTGTGTGAAGTGTGCGGCCAGGCCAGCGATCCTTCT<br>AGACTGCTGCTGTGCGACGACTGCGACATCAGCTACCAC<br>ACCTACTGTCTGGACCCTCCACTGCTGACCGTGCCTAAAG<br>GCGGATGGAAGTGCAAGTGGTGCGTGTCCTGTATGCAGT<br>GCGGAGCCGCCTCTCCTGGATTTCACTGCGAGTGGCAGA<br>ACAGCTACACCCACTGTGGCCCTTGTGCCAGCCTGGTCA<br>CCTGTCCTATTTGTCACGCCCCTTACGTGGAAGAGGACCT<br>GCTGATCCAGTGCAGACACTGCGAGAGATGGATGCACGC<br>CGGCTGCGAGAGCCTGTTCACAGAGGATGATGTGGAACA<br>GGCCGCCGACGAGGGCTTCGATTGTGTGTCTTGTCAGCC<br>CTACGTGGTCAAGCCCGTGGCTCCTGTTGCTCCTCCTGAA<br>CTTTAAGGATCCCTCGAGGCGGCCGCCCGCTGAGCAATA<br>ACTAGCA | ThermoScientific | 2D46_cellular |
| TACCTCACCGGTAGGCATATGTGTACAAGAGGAATGTCG<br>ACGAGATGGGCGGCCTGGGCATCCTGACATCTGTGCCCA<br>TCACTCCCAGAGTCGTGTGCTTCCTGTGTGCCTCTTCTGG<br>CCACGTGGAATTCGTGTACTGCCAAGTGTGCTGCGAGCC<br>CTTCCAGAAGTTCTGCCTGGAAGAGAACGAGCGGCCCCT | ThermoScientific | 2A13* H1456Q |

| | | |
|---|---|---|
| GGAAGATCAGCTGGAAAACTGGTGCTGCCGGCGGTGCAA GTTCTGTCATGTCTGTGGCAGACAGCACCAGGCCACCAA ACAGCTGCTGGAATGCAACAAGTGCCGGAACAGCTATCA CCCCGAGTGTCTGGGCCCTAACTACCCCACCAAGCCTAC CAAGAAAAAGAAAGTGTGGATCTGCACCAAATGCGTCCG GTGCAAGAGCTGCGGCTCTACCACACCTGGCAAAGGCTG GGATGCTCAGTGGTCCCACGATTTCAGCCTGTGCCACGA TTGCGCCAAGCTGTTCGCCAAGGGCAACTTCTGCCCTCTG TGCGACAAGTGCTACGACGACGATGACTACGAGAGCAA GATGATGCAGTGCGGCAAGTGCGACAGATGGGTGCACTC CAAGTGCGAGAATCTGAGCGGCACCGAGGACGAGATGT ACGAGATCCTGAGCAACCTGCCTGAGAGCGTGGCCTACA CCTGTGTGAACTGCACCGAAAGACACCCTGCCGAATGGC GGCTGGCCCTGGAAAAAGAACTTTAAGGATCCCTCGAGG CGGCCGCCCGCTGAGCAATAACTAGCA | | |
| TACCTCACCGGTAGGCATATGTGTACAAGAGGAATGTCG ACGAGGAAGAAGAAGAGGACGACGATACCATGCAGAAC ACCGTGGTGCTGTTCAGCAACACCGACAAGTTCGTGCTG ATGCAGGATATGTGCGTCGTGTGCGGCAGCTTTGGAAGA GGCGCTGAAGGACATCTGCTGGCCTGTAGCCAGTGCTCC CAGTGCTATTACCCCTACTGCGTGAACAGCAAGATCACC AAAGTGATGCTGCTGAAAGGCTGGCGCTGCGTGGAATGC ATCGTGTGTGAAGTGTGCGGCCAGGCCAGCGATCCTTCT AGACTGCTGCTGTGCGACGACTGCGACATCAGCTACCAC ACCTACTGTCTGGACCCTCCACTGCTGACCGTGCCTAAAG GCGGATGGAAGTGCAAGTGGTGCGTGTCCTGTATGCAGT GCGGAGCCGCCTCTCCTGGATTTCACTGCGAGTGGCAGA ACAGCTACACCCACTGTGGCCCTTGTGCCAGCCTGGTCA CCTGTCCTATTTGTCACGCCCCTTACGTGGAAGAGGACCT GCTGATCCAGTGCAGACACTGCGAGAGATGGATGCACGC CGGCTGCGAGAGCCTGTTCACAGAGGATGATGTGGAACA GGCCGCCGACGAGGGCTTCGATTGTGTGTCTTGTCAGCC CTACGTGGTCAAGCCCGTGGCTCCTGTTGCTCCTCCTGAA CTTTAAGGATCCCTCGAGGCGGCCGCCCGCTGAGCAATA ACTAGCA | ThermoScientifc | 2D46* H1405Y |
| TACCTCACCGGTAGGCATATGTGTACAAGAGGAATGTCG ACAAGAAAGGCCGCAGATCTCGGAGATGCGGCCAGTGTC CTGGATGCCAGGTTCCAGAAGATTGCGGCGTCTGCACCA ACTGCCTGGACAAGCCTAAGTTCGGCGGCAGAAACATCA AGAAACAGTGCTGCAAGATGCGGAAGTGCCAGAACCTG CAGTGGATGCCTTCTAAGGGCTCTGCCGGATCTGCTGCC GGAAGCGGAGAGTTTGAAATGGGCGGACTGGGCATCCTG ACCAGCGTGCCAATCACTCCTAGAGTCGTGTGCTTCCTGT GTGCCAGCTCTGGCCACGTGGAATTCGTGTACTGCCAAG TGTGCTGCGAGCCCTTCCACAAGTTCTGCCTGGAAGAGA ACGAGCGGCCCCTGGAAGATCAGCTGGAAAACTGGTGCT GCCGGCGGTGCAAGTTCTGTCATGTCTGTGGCAGACAGC ACCAGGCCACCAAACAGCTGCTGGAATGCAACAAGTGCC GGAACAGCTATCACCCCGAGTGTCTGGGCCCTAACTACC CCACCAAGCCTACCAAGAAAAAGAAAGTGTGGATCTGCA CTAAGTGCGTCCGGTGCAAGAGCTGCGGCTCTACCACAC CTGGCAAAGGCTGGGATGCTCAGTGGTCCCACGATTTCA GCCTGTGCCACGATTGCGCCAAGCTGTTCGCCAAGGGCA ACTTCTGCCCTCTGTGCGACAAGTGCTACGACGACGATG ACTACGAGAGCAAGATGATGCAGTGCGGCAAGTGCGAC AGATGGGTGCACTCCAAGTGCGAGAATCTGAGCGACGAG ATGTACGAGATCCTGAGCAACCTGCCTGAGAGCGTGGCC TACACCTGTGTGAACTGCACCGAAAGACACCCTGCCGAA TGGCGGCTGGCCCTGGAAAAAGAACTTTAAGGATCCTCT AGATAACTGAATTCTCGACCTCCGCCCGCTGAGCAATAA | ThermoScientific | 2ACP |

| | | |
|---|---|---|
| CTAGCAG | | |
| TACCTCACCGGTAGGCATATGTGTACAAGAGGAATGTCG ACGAGGAACCCCTGCTGGTCAACGTGGACAAAGCCGTGG TGTCTGGCTCTACCGAGAGATGCGCCTTCTGCAAACACCT GGGCGCCACCATCAAGTGCTGCGAAGAGAAGTGCACCCA GATGTATCACTACCCCTGTGCCGCTGGCGCCGGAACCTTT CAGGATTTCAGCCACATCTTTCTGCTGTGCCCCGAGCACA TCGATCAGGCCCCTGAGAGAAGCAAAGAGGACGCCAATT GCGCCGTGTGCGATAGCCCTGGCGATCTGCTGGATCAGT TCTTCTGCACCACCTGTGGCCAGCACTACCACGGCATGTG CCTGGATATTGCCGTGACACCCCTGAAGCGAGCCGGATG GCAATGCCCTGAGTGCAAAGTGTGCCAGAACTGCAAGCA GAGCGGCGAGGACAGCAAGATGCTCGTGTGCGACACCTG TGACAAGGGCTACCACACCTTCTGCCTGCAACCTGTGAT GAAGTCCGTGCCTACCAACGGCTGGAAGTGCAAGAACTG CCGGATCTGCATCGAGTGCGGCACCAGATCTAGCAGCCA GTGGCACCACAATTGCCTGATCTGCGACAACTGCTACCA GCAGCAGGACAATCTGTGCCCCTTCTGCGGCAAGTGCTA TCACCCCGAGCTGCAGAAAGACATGCTGCACTGCAACAT GTGCAAGAGATGGGTGCACCTGGAATGCGACAAGCCCAC CGATCACGAGCTGGACACCCAGCTGAAAGAAGAGTACAT CTGCATGTATTGCAAGCACCTCGGAGCCGAGATGGACAG ACTGCAGCCTGGCGAAGAAGGATCTGCAGGCTCTGCTGC AGGCAGCGGCGAGTTCGACGATGAAGAGAACAGCATGC ACAACACCGTGGTGCTGTTCAGCAGCTCCGACAAGTTCA CCCTGAACCAGGATATGTGCGTCGTGTGCGGCTCTTTTGG ACAGGGCGCTGAAGGCAGACTGCTGGCCTGTTCTCAGTG CGGACAGTGCTACCATCCTTACTGCGTGTCCATCAAGATC ACCAAGGTGGTGCTGAGCAAAGGCTGGCGGTGCCTGGAA TGTACCGTGTGTGAAGCCTGTGGCAAGGCCACAGATCCT GGAAGGCTGCTGCTGTGCGACGACTGCGATATCAGCTAT CACACCTACTGTCTGGACCCTCCACTGCAGACCGTGCCTA AAGGCGGATGGAAATGCAAGTGGTGCGTGTGGTGTAGAC ACTGCGGCGCTACATCTGCCGGACTGAGATGCGAGTGGC AGAACAACTACACCCAGTGCGCCCCTTGTGCCAGCCTGA GTTCTTGCCCTGTGTGCTACCGGAACTACAGAGAAGAGG ACCTGATCCTGCAGTGCCGGCAGTGCGATAGATGGATGC ATGCCGTGTGTCAGAACCTGAACACCGAGGAAGAGGTGG AAAACGTCGCCGACATCGGCTTCGACTGCTCCATGTGCA GACCCTACATGCCCGCCAGCAATGTGCCTAGCTCTGACT AAGGATCCTCTAGATAACTGAATTCTCGACCTCCGCCCG CTGAGCAATAACTAGCAG | ThermoScientific | 2C2P |
| TACCTCACCGGTAGGCATATGTGTACAAGAGGAATGTCG ACAGCAAGCAGTTCGTGCACAGCAAGAGCAGCCAGTACC GGAAGATGAAGACCGAGTGGAAGTCCAACGTGTACCTG GCCAGAAGCAGAATCCAAGGCCTGGGCCTGTATGCCGCC AGAGACATCGAGAAGCACACCATGGTCATCGAGTACATC GGCACCATCATCCGGAACGAGGTGGCCAACCGGAAAGA GAAGCTGTACGAGAGCCAGAACCGGGGCGTGTACATGTT CAGAATGGACAACGACCACGTGATCGACGCCACACTGAC AGGTGGCCCTGCCAGATATATCGCCCACAGCTGCGCCCC TAATTGCGTGGCCGAGGTGGTCACATTTGAGCGGGGCCA CAAGATCATCATCAGCAGCTCCAGAAGAATCCAGAAAGG CGAGGAACTGTGCTACGACTACAAGTTCGACTTCGAGGA CGACCAGCACAAGATCCCTTGTCACTGTGGCGCCGTGAA CTGCCGGAAGTGGATGAACTAAGGATCCTCTAGATAACT GAATTCTCGACCTCCGCCCGCTGAGCAATAACTAGCAG | ThermoScientific | 2CS |
| TACCTCACCGGTAGGCATATGTGTACAAGAGGAATGTCG ACGAGCTGTGCGGCGTGGACAAGGCCATCTTTAGCGGCA TCAGCCAGAGATGCAGCCACTGCACAAGACTGGGCGCCA | ThermoScientific | 2D2P |

| | | |
|---|---|---|
| GCATTCCTTGTAGAAGCCCTGGCTGCCCCAGACTGTATCA<br>CTTCCCTTGTGCTACAGCCAGCGGCAGCTTCCTGAGCATG<br>AAGACACTGCAGCTGCTGTGCCCTGAGCACTCTGAAGGC<br>GCCGCTTATCTGGAAGAGGCCAGATGCGCTGTGTGTGAA<br>GGACCTGGCGAGCTGTGTGACCTGTTCTTCTGTACCAGCT<br>GCGGCCACCACTATCACGGCGCCTGTCTGGATACAGCCC<br>TGACCGCCAGAAAACGCGCTGGATGGCAATGCCCCGAGT<br>GCAAAGTTTGCCAGGCCTGCAGAAAGCCCGGCAACGACT<br>CTAAGATGCTCGTGTGCGAGACATGCGACAAGGGCTACC<br>ACACCTTCTGCCTGAAGCCTCCAATGGAAGAACTGCCCG<br>CTCACAGCTGGAAGTGCAAGGCCTGTAGAGTGTGCAGAG<br>CCTGTGGCGCCGGATCTGCCGAGCTGAATCCTAACTCTG<br>AGTGGTTCGAGGGCAGCGCCGGTTCTGCTGCTGGATCTG<br>GCGAGTTCGAGGAAGAAGAAGAGGACGACGATACCATG<br>CAGAACACCGTGGTGCTGTTCAGCAACACCGACAAGTTC<br>GTGCTGATGCAGGATATGTGCGTCGTGTGCGGCTCCTTTG<br>GCAGAGGCGCTGAAGGACATCTGCTGGCCTGTAGCCAGT<br>GCTCCCAGTGCTATCACCCCTACTGCGTGAACAGCAAGA<br>TCACCAAAGTGATGCTGCTGAAAGGCTGGCGCTGCGTGG<br>AATGCATTGTGTGCGAAGTGTGCGGCCAGGCCAGCGATC<br>CTAGTAGACTGCTGCTGTGTGATGACTGCGACATCTCCTA<br>CCACACATACTGTCTGGACCCTCCACTGCTGACCGTGCCT<br>AAAGGCGGATGGAAATGCAAGTGGTGCGTGTCCTGTATG<br>CAGTGCGGAGCCGCCTCTCCTGGATTTCACTGCGAGTGG<br>CAGAACAGCTACACCCACTGTGGCCCTTGTGCCAGCCTG<br>GTCACCTGTCCTATTTGTCACGCCCCTTACGTGGAAGAGG<br>ACCTGCTGATCCAGTGCAGACACTGCGAGAGATGGATGC<br>ACGCCGGCTGCGAGAGCCTGTTCACAGAGGATGATGTGG<br>AACAGGCCGCCGACGAGGGCTTCGATTGTGTGTCTTGTC<br>AGCCCTACGTGGTCAAGCCCGTGGCTCCTGTTGCTCCTCC<br>TGAACTTTAAGGATCCTCTAGATAACTGAATTCTCGACCT<br>CCGCCCGCTGAGCAATAACTAGCAG | | |
| TACCTCACCGGTAGGCATATGTGTACAAGAGGAATGTCG<br>ACGAGACAAACACCCCTTACAGCAAGCAGTTCGTGCACA<br>GCAAGAGCAGCCAGTACAGACGGCTGAGAACCGAGTGG<br>AAGAACAACGTGTACCTGGCCAGAAGCAGAATCCAAGG<br>CCTGGGACTGTACGCCGCCAAGGACCTGGAAAAGCACAC<br>CATGGTCATCGAGTACATCGGCACCATCATCCGGAACGA<br>GGTGGCCAACAGAAGAGAGAAGATCTACGAGGAACAGA<br>ACCGGGGCATCTACATGTTCCGGATCAACAACGAGCACG<br>TGATCGACGCCACACTGACAGGTGGCCCTGCCAGATATA<br>TCGCCCACAGCTGCGCCCCTAATTGCGTGGCCGAGGTGG<br>TCACCTTCGACAAAGAGGACAAGATCATCATCATCTCCA<br>GCCGGCGCATCCCCAAGGGCGAAGAACTGACCTACGACT<br>ACCAGTTCGACTTCGAGGACGACCAGCACAAGATCCCTT<br>GTCACTGCGGCGCCTGGAACTGCCGGAAATGGATGAACT<br>AAGGATCCTCTAGATAACTGAATTCTCGACCTCCGCCCG<br>CTGAGCAATAACTAGCAG | ThermoScientific | 2DS |

**Tabel S2. Biotinylated peptides used in this study.**

| PEPTIDE SEQUENCE | SOURCE | IDENTIFIER |
|---|---|---|
| ARTKQTARKSTGGKAPRKQLATKAARKSAP | GenScript | H3 |
| ARTKQTAR**Kme3**STGGKAPRKQLATKAARSAP | GenScript | H3K4me3 |
| ARTKQTAR**Kac**STGGKAPRKQLATKAAR**Kac**SAP | GenScript | H3K9acK27ac |

**Tabel S3. Oligonucleotides used in this study.**

| OLIGONUCLEOTIDE SEQUENCE | SOURCE | IDENTIFIER |
|---|---|---|
| **Cloning using SLIC to pEYFP** | | |
| CGGTAGGCATATGTGTACAAGAG | Genomed | Synthetic_F |
| TAGTTATTGCTCAGCGGGCG | Genomed | Synthetic_R |
| GCGGCTGGCCCTGGAAAAAGAACTT TGAGGATCCACCGGATCTAGAT | Genomed | PEY_A13_F |
| TCAGGATGCCCAGGCCGCCCATCTC GTCGACTGCAGAATTCGAAGCT | Genomed | PEY_A13_R |
| GGACAGACTGCAGCCTGGCGAAGAA TGAGGATCCACCGGATCTAGAT | Genomed | PEYFP_C14_F |
| TGTCCACGTTGACCAGCAGGGGCTC GTCGACTGCAGAATTCGAAGCT | Genomed | PEYFP_C14_R |
| TGCTGTTCTCTTCGTCGTCGTCGACTGCAGAATTCGAAG | Genomed | PEY_C57-CELL-OPTR |
| ATGTGCCTAGCTCTGACTAAGGATCCACCGGATCTAGA TAACTG | Genomed | PEY_C57-CELL-OPTF |
| TGGCCTTGTCCACGCCGCACAGCTCGTCGACTGCAGAA TTCGAAG | Genomed | PEY_D13-CELL-OPTR |
| GAATCCTAACAGCGAGTGGTTCGAATGAGGATCCACCG GATCTAGAT | Genomed | PEY_D13-CELL-OPTF |
| TGCTCCTCCTGAACTTTAAGGATCCACCGGATCTAG | Genomed | PEY_D46-CELL-OPTF |
| GGTATCGTCGTCCTCTTCTTCTTCCTCGTCGACTGCAGA ATTCGAAG | Genomed | PEY_D46-CELL-OPTR |
| **Cloning using SLIC from pEYFP to pLJM1** | | |
| CCATTTGTCTCGAGGTCGAGAATTCAGTTATCTAGATCC GGTGGATCC | Sigma-Aldrich | revSLICpljm1 |
| CGTCAGATCCGCTAGCGCTACC | Sigma-Aldrich | 5UTRpCMV |

**Table S4. Key reagent used in this study.**

| REAGENT | SOURCE | Catalog numer |
|---|---|---|
| **Antibodies** | | |
| anti-GFP-Mnase | Nizamuddin et al, 2021; kind gift from Marc Timmers | N/A |
| Anti-GFP | R&D Systems | Cat#AF4240 |
| anti-GST-HRP | Abcam | Cat#ab3416 |
| anti-H3 | Cell Signaling Technology | Cat#4499 |
| normal rabbit IgG | Sigma-Aldrich | Cat#I8140 |
| anti-KMT2A | Diagenode | Cat#C15310264 |
| anti-KMT2D | Invitrogen | Cat#701869 |
| anti-WDR5 | Diagenode | Cat#C15410027 |
| anti-RbBP5 | ActiveMotif | Cat#61406 |
| anti-H3K4me1 | Cell Signaling Technology | Cat#5326 |
| anti-H3K4me3 | Cell Signaling Technology | Cat#9751 |
| anti-H3K9ac | Diagenode | Cat# C15410004 |
| anti-H3K27ac | Diagenode | Cat# C15410196 |
| anti-H3K23ac | Diagenode | Cat# C15410344 |
| anti-H3K56ac | Diagenode | Cat# C15410213 |
| anti-H2A.Zac | Diagenode | Cat#C15410173 |

| | | |
|---|---|---|
| anti-H2BK15ac | Diagenode | Cat# C15410220 |
| anti-H4K16ac | ActiveMotif | Cat#39168 |
| anti-H4K5acK8acK12ac | Diagenode | Cat#C15410021 |
| anti-H4K5acK8acK12acK16ac | Diagenode | Cat#C15410024 |
| **Chemicals and recombinant proteins** | | |
| MNase | New England Biolabs | Cat#M0247S |
| CUT&RUN pAG-MNase | Cell Signaling Technologies | Cat#40366 |
| Digitonin | Sigma-Aldrich | Cat#D141 |
| Spermatidine | Sigma-Aldrich | Cat#S2626 |
| BioMag®Plus Concanavalin A | Polysciences | Cat# 86057-3 |
| GFP-Trap magnetic Agarose | Chromotek/Proteintech | Cat# gtma |
| Dynabeads MyOne Streptavidin C1 | ThermoScientific | Cat#65001 |
| cOmplete mini EDTA-free protease inhibitors cocktail | Sigma-Aldrich | Cat#4693159001 |
| Recombinant protein H4 | New England Biolabs | Cat#M2504S |
| Recombinant protein H3.1 | New England Biolabs | Cat#M2503S |
| ChIP DNA Clean & Concentrator™ | Zymo Research | Cat#D5201 |
| NEBNext® Ultra™ II DNA Library Prep Kit for Illumina® | New England Biolabs | Cat#E7645 |
| NEBNext® Multiplex Oligos for Illumina® (Index Primers Set 1) | New England Biolabs | Cat#E7335 |
| NEBNext® Multiplex Oligos for Illumina® (Index Primers Set 2) | New England Biolabs | Cat#E7500 |
| NEBNext® Multiplex Oligos for Illumina® (Index Primers Set 3) | New England Biolabs | Cat#E7710 |
| NEBNext® Multiplex Oligos for Illumina® (Index Primers Set 4) | New England Biolabs | Cat#E7730 |
| NovaSeq 6000 S1 Reagent Kit (200 cycles) | Illumina | Cat#20028318 |
